# A computational model for investigating the evolution of colonic crypts during Lynch syndrome carcinogenesis

**DOI:** 10.1101/2020.12.29.424555

**Authors:** Saskia Haupt, Nils Gleim, Aysel Ahadova, Hendrik Bläker, Magnus von Knebel Doeberitz, Matthias Kloor, Vincent Heuveline

**Affiliations:** Engineering Mathematics and Computing Lab (EMCL), Interdisciplinary Center for Scientific Computing (IWR), Heidelberg University, Heidelberg, Germany; Data Mining and Uncertainty Quantification (DMQ), Heidelberg Institute for Theoretical Studies (HITS), Heidelberg, Germany; Department of Applied Tumor Biology (ATB), Institute of Pathology, University Hospital Heidelberg, Heidelberg, Germany; Clinical Cooperation Unit Applied Tumor Biology, German Cancer Research Center, Heidelberg, Germany; Institute of Pathology, University Hospital Leipzig, Leipzig, Germany

**Keywords:** Colonic Crypts, Colorectal Cancer, Carcinogenesis, Lynch Syndrome, Computational Modeling, Voronoi Tessellation Model, Monoclonal Conversion

## Abstract

Lynch syndrome (LS), the most common inherited colorectal cancer (CRC) syndrome, increases the cancer risk in affected individuals. LS is caused by pathogenic germline variants in one of the DNA mismatch repair (MMR) genes, complete inactivation of which causes numerous mutations in affected cells. As CRC is believed to originate in colonic crypts, understanding the intra-crypt dynamics caused by mutational processes is essential for a complete picture of LS CRC and may have significant implications for cancer prevention.

We propose a computational model describing the evolution of colonic crypts during LS carcinogenesis. Extending existing modeling approaches for the non-Lynch scenario, we incorporated MMR deficiency and implemented recent experimental data demonstrating that somatic *CTNNB1* mutations are common drivers of LS-associated CRCs, if affecting both alleles of the gene. Further, we simulated the effect of different mutations on the entire crypt, distinguishing non-transforming and transforming mutations.

As an example, we analyzed the spread of mutations in the genes *APC* and *CTNNB1*, which are frequently mutated in LS tumors, as well as of MMR deficiency itself. We quantified each mutation’s potential for monoclonal conversion and investigated the influence of the cell location and of stem cell dynamics on mutation spread.

The *in silico* experiments underline the importance of stem cell dynamics for the overall crypt evolution. Further, simulating different mutational processes is essential in LS since mutations without survival advantages (the MMR deficiency-inducing second hit) play a key role. The effect of other mutations can be simulated with the proposed model. Our results provide first mathematical clues towards more effective surveillance protocols for LS carriers.

## 1 | INTRODUCTION

Lynch syndrome (LS) is the most common hereditary colorectal cancer (CRC) syndrome affecting 1 in every 180 people ^1^. Besides an increased risk for many other types of cancer, LS leads to a lifetime risk of over 50% of developing CRC ^2^. LS is inherited via a pathogenic germline variant in one of the four mismatch-repair (MMR) genes *MLH1, MSH2, MSH6, PMS2* ^3^. A second somatic hit affecting the remaining functional allele of the same MMR gene leads to DNA MMR deficiency ^4^, resulting in the accumulation of insertion/deletion mutations, especially at repetitive DNA sequences (microsatellites). Consequently, cancers developing in LS show the molecular phenotype of microsatellite instability (MSI), i.e., length alterations of microsatellites. Mutations at microsatellites residing in gene-encoding regions may result in the inactivation of tumor suppressor genes and thereby drive carcinogenesis. Thus, MMR deficiency is a major driving force in LS carcinogenesis.

The epithelium of the colon is organized in crypts. Crypt epithelial cells are considered cells of origin for CRC ^5^. Proliferation is ongoing in stem cells and multiple times in transit-amplifying cells near the crypt base giving rise to mutations in both types of cells ^6;7;8^. The mutations can then spread throughout the crypt. A more comprehensive introduction into the biological details of cell dynamics within crypts is given in the Appendix A.

However, a detailed understanding of the intra-crypt dynamics underlying both healthy and aberrant crypts is still lacking. A reason for this is that *in vivo* experiments with a temporal resolution are very hard to implement in practice. In other words, there is a gap in understanding how a mutation in a single cell takes over a colonic crypt and how this contributes to Lynch syndrome carcinogenesis and thus results in tumor risk predictions on a population level.

In LS, it was possible to identify a unique precursor lesion of carcinomas, called MMR-deficient crypt foci. They are characterized by the loss of MMR protein expression in the absence of histological signs of dysplasia ^9^. MMR-deficient crypts are extremely hard to detect by conventional endoscopy approaches commonly applied in LS individuals and may therefore explain the frequently observed phenomenon of cancer manifestation despite recent colonoscopy ^10;11;12^. Therefore, understanding tumor development from these lesions is essential for designing tailored prevention strategies and reducing CRC incidence in LS (see Figure 1).

**FIGURE 1.**
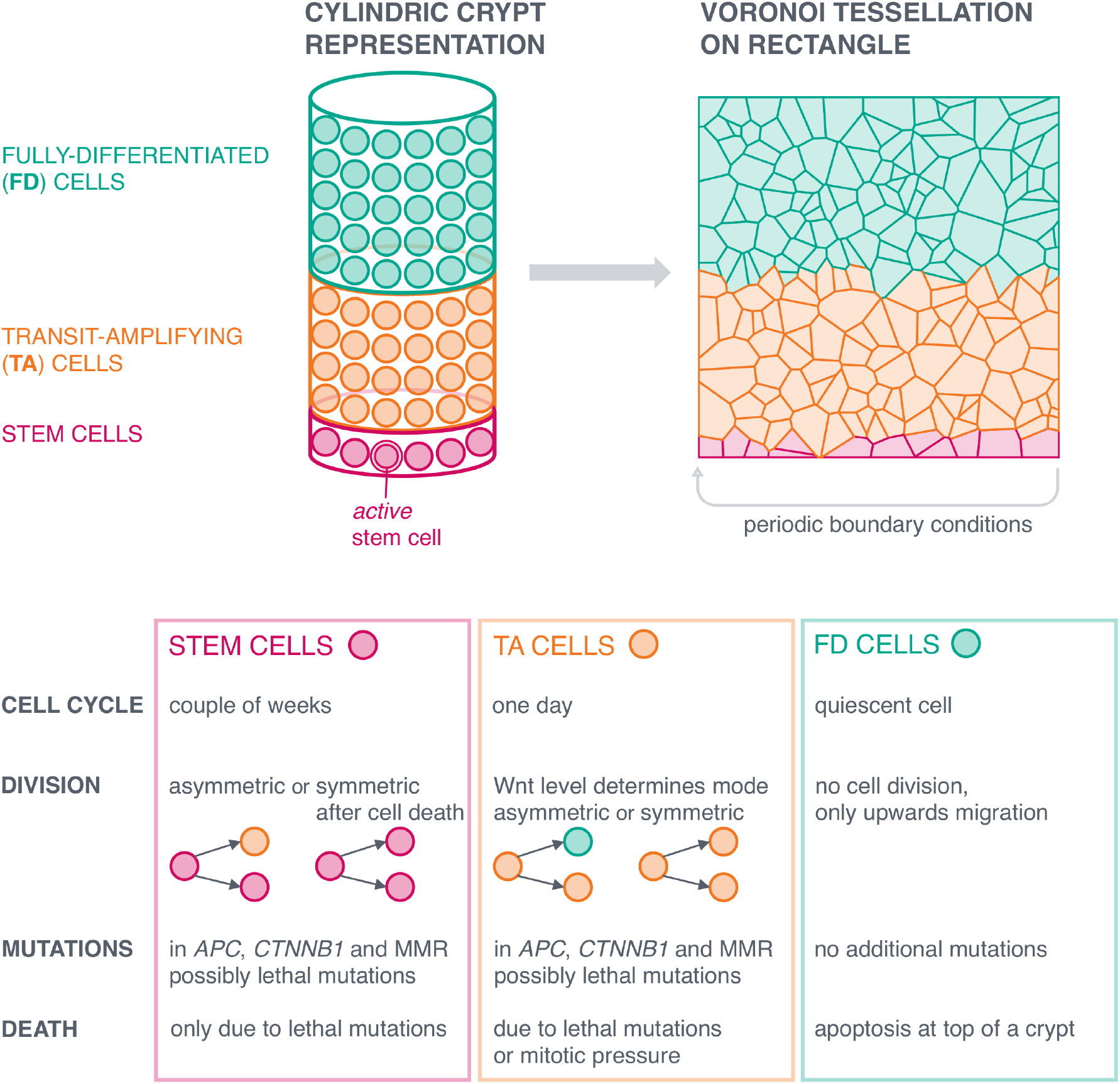
Overview of the computational model of colonic crypts. *Top:* The colonic crypt is represented by a cylinder consisting of stem cells (red) at the bottom, transit-amplifying cells (orange) in the middle and fully-differentiated (FD) cells (green) at the top of the crypt. An active stem cell populates the crypt at any point in time. As we model LS, all cells are initialized with a germline variant in exactly one of the MMR genes. The cylinder is transformed into a rectangle with periodic boundary conditions, where the cells are represented by a Voronoi tessellation. *Bottom:* For each cell type, we model the cell cycle including cell division, possible mutations in one of the MMR genes, in *APC* and *CTNNB1*, and multiple death mechanisms.

### 1.1 | Contribution

In this work, we present a computational model with numerical simulations of colonic crypt evolution during LS carcinogenesis (see Figure 1). The main goal of the modeling approach is to translate knowledge about the effects of defined mutations from the cellular to the crypt level. Although experimental data on mutation rates in dividing cells *in vitro* are existing, it is hard to translate these numbers onto the level of crypts, the organ, or the individual. In these lines, information about (1) the likelihood of a defined mutation leading to monoclonal conversion of the surrounding crypt, and (2) the time until conversion takes place are paramount. The present model has been designed as a first step to fill this gap of knowledge.

Although organoid cultures represent a huge leap forward in the analysis of cellular alterations in a tissue context, computational models are more flexible with regard to altering certain parameters, including the implementation of environmental changes and their effects on crypt homeostasis or mutational manifestation. Moreover, computational models have the significant advantage that upscaling the number of simulations is only limited by the availability of computational resources.

The present model is an extension and adaptation of existing approaches ^13;14;15^ for modeling LS carcinogenesis allowing to obtain *in silico* experiments for mutational processes and intra-crypt dynamics during LS carcinogenesis. We take into account crucial biological features by defining stem cell dynamics with only one active stem cell at a time according to ^16;17^. In addition, the modeling incorporates feedback and resulting death mechanisms which turn out to be essential to avoid an overpopulation of the crypt.

Besides somatic *APC* mutations as known drivers of CRC in general, we incorporate MMR deficiency and related genetic dependencies such as increased mutation and death rates into the model. Further, we make use of recent experimental data ^18^ demonstrating that somatic *CTNNB1* mutations, if affecting both alleles of the gene, are common drivers of LS-associated CRCs.

Our model is able to simulate the effect of different mutations, including non-transforming and transforming mutations, on the intra-crypt dynamics. Mutations without any survival advantage, such as the MMR deficiency-inducing second hit are of particular importance in LS carcinogenesis. Modeling the effects of both types of mutations is thus essential for the biological understanding. We investigate how these driver mutations influence the intra-crypt dynamics and analyze the influence of the cell location and the effect of stem cell dynamics on the spread and monoclonal conversion of mutations within a crypt. Effects of other, possibly yet unidentified mutations of both non-transforming and transforming type can be simulated with the proposed model.

The model parameters are based on the existing biomedical and clinical estimates allowing a comparison with available human data and new medical hypotheses for human colonic crypt evolution. Besides that, it is possible to use the modeling approach for murine colonic crypts by adapting size- and species-dependent parameters, which can support studies analyzing carcinogenic processes using animal models. Further, upon adaptation of certain parameters, this model can be rolled-out to reflect the development of sporadic CRCs and CRCs in the Lynch-like ^19^ and familial adenomatous polyposis (FAP) scenario.

### 1.2 | Related Work

The mathematical approaches used to model cell populations can be broadly divided into three categories: 1) *Spatial models*, which take into account the specific location of individual cells or the location of a population of cells, 2) *compartmental models*, which describe the transition between cell types, irrespective of their position within the population, and 3) non-spatial *stochastic models*, a more general class of models, all involving stochasticity as a main feature. For detailed reviews, we refer to ^20;7;21;22;23;24^. It is important to note that the models are distinguished based on their basic setup, not on the mathematical tools they use. For instance, one can couple a setup with ordinary (ODEs) or partial differential equations (PDEs) as well as stochastic processes. This gives rise to a *system*, whereby the methods are used to describe the *state* of the system.

As our model is a spatial model, we will explain the main ideas of such an approach following the reviews mentioned above. We would like to point out that we are referring to cell-based spatial models in the following using a discrete spatial representation of cells rather than a continuous representation by, e.g., PDE-based approaches.

**Cell-based spatial models** treat cells as discrete entities bearing specific characteristics as internal states which change in discrete time steps. At each time step, these internal states are updated according to certain rules (usually governed by equations), such that the whole state of the system is recomputed. Cell-based spatial models can be further divided into two subcategories: *in-lattice models* and *off-lattice models*. The main structural difference lies in whether or not the cells are assumed to be positioned on a rigid grid.

In off-lattice models, cells are loosely located in space, yielding cell shapes which are more biologically realistic. Models within this class are further distinguished depending on how exactly cells are labeled within the underlying space, such that we can track their position over time. *Vertex models* represent cells as polygons and the vertices of the cells are tracked in space and time, while *overlapping spheres (OS) models* and *Voronoi tessellation (VT) models* describe cells through the position of their nuclei and the nuclei are tracked in space and time. OS models regard cells as spheres with a certain time-dependent radius. Voronoi tessellations give rise to a cell body consisting of all points in space whose distance to the nucleus is less than or equal to their distance to any other nucleus in the cell population. Each model is equipped with a unique representation of cell division. VT models have been used extensively to model cellular migration ^13^, or in multi-scale models ^25^, which used VT to model the tissue architecture of stratified epithelium. We will make use of the VT model for cellular migration ^13^ in our computational model for the realization of cell migration. Not all modeling approaches which concern intestinal crypts can clearly be assigned to exactly one of the three major classes. Many models concentrate on single aspects rather than cell populations and can further be used in multi-scale models.

There are two approaches, which investigate similar aspects as our study ^15;26;27^. Both focus on the process of monoclonal conversion (also called mutation fixation) in intestinal crypts, using multi-scale models based on spatial approaches. The first uses Voronoi tessellations and a similar cell migration realization, also based on the Meineke model ^13^, but represents the crypt as a two-dimensional surface of revolution rather than a cylinder. Further, this model shares assumptions with our model regarding the role of the Wnt pathway for differentiation. However, the representation of stem cell dynamics, as well as assumptions on crypt size and cell cycle lengths differ substantially from ours, rendering the model more suitable for application to mouse data than to human data. This feature also concerns the approach by Araujo et al., which is based on a cellular automaton model, an example of an in-lattice model, rather than an off-lattice model, which entails limitations regarding the cells’ geometric representation. Gene-gene interactions are incorporated in more detail compared to our approach, via the use of a gene regulatory network, however the implementation of sporadic mutations is lacking.

Regarding the modeling of the Wnt pathway, the approaches in ^28;20^ share our assumption that the Wnt level is one of the primary regulators of cell differentiation (see also Appendix A). Further, ^28^ take into account that the colonic crypt has a cap, and by this, assume that Wnt activity is determined by the local curvature of the basal membrane. However, we use a rather simplified cylindric geometry to reduce complexity and computational costs.

## 2 | MODELING CRYPT EVOLUTION

We present a computational model for the crypt dynamics based on a Voronoi tessellation ansatz with parameters set according to published data (see Appendix B). We assume that a crypt can be geometrically represented by an open cylinder. This is implemented by considering a two-dimensional rectangular surface Ω = [0, 2*πr*_crypt_] × [0, *h*_crypt_] with surface area *L*_Ω_ = 2*πr*_crypt_*h*_crypt_ and periodic boundary conditions. We note that this geometric representation constitutes a simplification, and that other approaches as used in ^28;15^ provide extensions of our approach.

For the intra-crypt dynamics, we assume a Voronoi tessellation model, which has been shown to be a well-suited approach to model epithelia ^29^. By this means, the body of the cell is given by the Voronoi tessellation using Euclidean distances (see Figure 1, upper right).

### Definition 1 (Voronoi Tessellation)

*Consider a population of k* ∈ ℕ *cells denoted as the set* {1, …, *k*}, *together with the positions of their nuclei* (*r*_*i*_)_*i* =1,…,*k*_ ⊆ Ω. *The* Voronoi cell *(cell body) is then defined as*

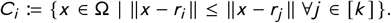

*The collection of cells* (*C*_*i*_)_*i* =1,…,*k*_ *is called* Voronoi tessellation.

The term *cell nucleus* in our model does not refer to a physical subcellular compartment or structure but is defined as the geometric center of the Voronoi tessellation. It also has a biological meaning with regards to the modeling of division by placing the defined nuclei of cells after division at a fixed distance to each other (see Section 2.3).

### 2.1 | Modeling the Cell Cycle

For each cell, we consider a cell cycle model to regulate cell proliferation, and to determine circumstances for cell differentiation and mutations. The cell cycle consists of different phases (see Appendix A) which are all implemented in our model. The duration of each cell cycle is assumed to vary stochastically around a mean length for different cells. While the other phases are assumed to be fixed, the G1 phase varies from cell to cell in our model. Experimentally, the exact distribution is hard to determine ^30^, and we assume that the G1 phase follows a normal distribution *t* _G1_ ∼ 𝒩 (*m*_G1_, *σ* _G1_) ^30^. The values for *m*_G1_ and *σ* _G1_ may vary for different cell types.

In the cell cycle model, the activity of the Wnt pathway distinguishes the dividing cells, i.e., stem cell and TA cells, from the non-dividing FD cells. In other words, the activity of the Wnt pathway determines the cell type. However, we assume only the active stem cell to be responsive to Wnt signaling, such that the non-dividing quiescent stem cells are non-responsive. Thus, we assume that each cell is assigned a Wnt level *l*_wnt_, where the lower the cell is located in the crypt, the higher is its Wnt level.

#### Definition 2 (Wnt Level)

*The Wnt level l*_wnt_ *is defined by*

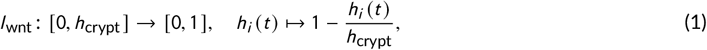

*where h*_crypt_ ∈ ℝ_>0_ *denotes the height of the crypt and the current height h*_*i*_ (*t*) *is given by the y-coordinate of the position of the cell nucleus*.

For distinguishing the cell types, we introduce a Wnt threshold *τ*_wnt_, where

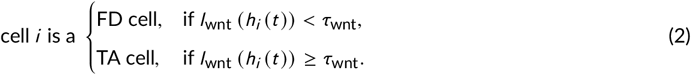

### 2.2 | Modeling Cell Differentiation

The modes of cell differentiation differ for the different cell types. For a summary, we refer to Figure 1. Cell differentiation for stem cells is assumed to be asymmetric, always leading to one stem cell and one TA cell. Only in the case that one stem cell dies, the neighboring stem cell divides symmetrically such that the total number of stem cells is fixed in time. For TA cells, the Wnt level introduced above determines the way of cell differentiation leading either to two TA cells or to one TA and one FD cell. FD cells are assumed to never divide, and cannot differentiate any further.

### 2.3 | Modeling Cell Division and Mutations

During cell division, the daughter cells are created. In Voronoi tessellations, this is done by placing the daughter cell nucleus *i*_*d*_ at a close, fixed distance *є* from the mother cell nucleus *i*_*m*_ in a random direction, i.e., ‖*i*_*d*_ − *i*_*m*_ ‖ = *є*. Subsequently, the tessellation is recomputed. We note that the random placement of daughter cell nuclei in all directions in our model is a more general approach of cell division compared to the fixed lateral placement to the right or to the left in ^31^. As there seems to be no biological evidence for a restriction of daughter nucleus placement, we have chosen the more general approach in this respect.

The cell cycle length *t*_*cc*_ (*i*_*m*_) of *i*_*m*_ is fixed, whereas a new cell cycle length is assigned to *i*_*d*_ with the G1 phase length sampled from 𝒩 (*m*_G1_, *σ* _G1_). The ages are set to 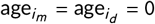 for both cells *i*_*m*_ and *i*_*d*_.

During each cell division, the new daughter cell can acquire mutations. The following modeling ansatz with the corresponding assumptions corresponds to the one presented in ^32^. We model mutational events independent of and dependent on other mutations and assume a gene-dependent point mutation rate *π*_mut_ (gene) and LOH event rate *p*_LOH_ (gene). More details are given in the Appendix B.

The genotypic state *g* (*i, t*) of a cell *i* at time *t* is determined by the mutation state of the three genes (one of the MMR genes, *APC* and *CTNNB1*) at this time given by the triple

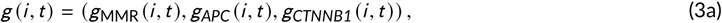

where for all cells *i*, for all time points *t*

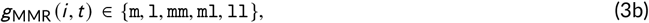

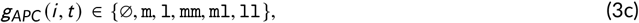

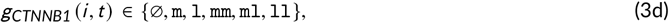

where ∅ denotes a wild-type gene, `m` denotes a point mutation, and `l` an LOH event. As we focus on the LS scenario, all cells have a first hit in the respective MMR gene, i.e., *g*_MMR_ (*i*, 0) = `m` ∀*i* or *g*_MMR_ (*i*, 0) = `l` ∀*i*. We neglect the possibility that two somatic mutations occur in one of the other MMR genes. Further, we assume that two LOH events in *APC* or *CTNNB1* damage the cell in such a way that it directly leads to cell death. According to ^33^, somatic *CTNNB1* mutations are significantly more frequent in *MLH1*-associated LS CRCs compared to CRCs associated with the other MMR genes. For illustration purposes, we make the assumption that inactivation of *MLH1* and *CTNNB1* are triggered by non-independent events. We incorporate this dependency in our model with an occurrence rate *r*_effLOH_. We assume that the second hit at time *t* _2_ in an MMR gene leads to an increased point mutation rate *p*_mut_ by a factor *λ*_mut_ > 1 for all other genes in this cell, i.e., if *g*_MMR_ (*i, t*) ∈ {`mm, ml, ll`}, then

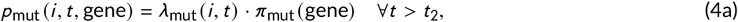

where

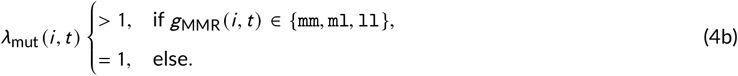

The exact parameter setting used for the simulations is given in Appendix B. Mutations in both *APC* and *CTNNB1* are assumed to increase the activity of the Wnt pathway. This is implemented in the model by decreasing the Wnt threshold *τ*_wnt_ by a predefined factor *λ*_wnt_ (*i, t*) ∈ [0, 1], which is equivalent to increasing the Wnt level for those cells. The factor varies for different genotypic states. It is assumed to be the smallest for biallelically *APC*- or *CTNNB1*-mutated cells, and larger for monoallelically *APC*-mutated and monoallelically *CTNNB1*-mutated cells. In formulas, this reads

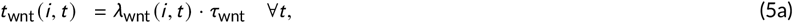

where

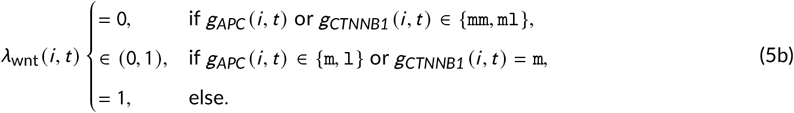

The exact parameter setting used for the simulations is given in Appendix B. Further, the effects of the different mutations on the intra-crypt dynamics are summarized in Table 1.

**TABLE 1.** Effects of mutations on intra-crypt dynamics.. We summarize the effects on mutation rates, differentiation and cell death for MMR deficiency, monoallelic and biallelic *APC* or *CTNNB1* mutations, which are implemented in the model.

| Mutation | Effect on mutation rate | Effect on differentiation | Effect on death |
| --- | --- | --- | --- |
| MMR deficiency | increase | none | increase of mutation-induced death |
| monoallelic APC or CTNNB1 | none | delay | none |
| biallelic APC or CTNNB1 | none | inhibition | partial apoptotic resistance, cell death if complete gene loss |

### 2.4 | Modeling Cell Migration

We assume the cells of the colonic crypt to exert pressure on each other. This, together with the movement of cells due to cell divisions, contributes to the upward movement along the vertical crypt axis. Rather than explicitly modeling a tissue-level pressure, the mitotic pressure is implicitly incorporated into a force-based cell mechanics model. The results in Paulus et al. ^34^ suggest that upward cell migration continues until up to 20% of its original value after complete mitotic inhibition, indicating that mitotic pressure is the main contributor to upward migration. We use a *linear spring force*, as defined in ^13^. Here, we make the following two assumptions:

1. The forces between the cells are modeled by a network of springs, where the latter are the vectors connecting one cell to the other.
2. Each force linearly depends on this vector.

#### Definition 3 (Spring)

*Let r*_*i*_ (*t*), *r*_*j*_ (*t*) ∈ Ω *denote the positions of the nuclei of two cells i and j*. *Then, the vector r*_*ij*_ (*t*) = *r*_*j*_ (*t*) − *r*_*i*_ (*t*) *is called* spring *from cell i to cell j at time t*.

#### Definition 4 (Resting Spring Length)

*We consider cells i and j at time t, for which cell i does not exert pressure on cell j*. *Then, the* resting spring length, *denoted by s*_*ij*_ (*t*), *corresponds to the distance between the cell nuclei and is the “natural length” of the spring between those two cells. It is defined in the following way*

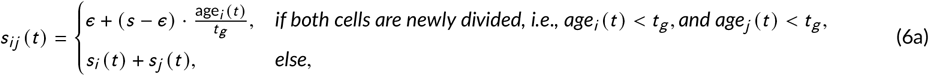

*where*

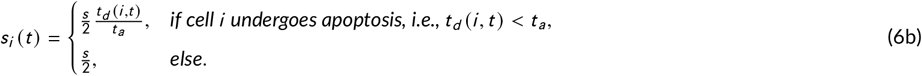

*Here, є* ∈ ℝ_>0_ *is the distance between two cell nuclei after division*, age_*i*_ (*t*) ∈ ℝ_≥ 0_ *is the age of cell i at time t, t*_*g*_ ∈ ℝ_>0_ *is the time needed for a newly divided cell to grow to its original size, t*_*d*_ (*i, t*) ∈ ℝ_≥ 0_ *is the time until the death of cell i, and t* _*a*_ ∈ ℝ_>0_ *the duration of apoptosis. Further, the parameters s* ∈ ℝ_>0_, *є, t*_*g*_ *and t* _*a*_ *are assumed to be time-independent and fixed for each cell*.

Directly after cell division, the distance between the two cells is *є*. After cell growth, this distance is increased to *s* ≥ *є*. As age_*i*_ (*t*) is linearly increasing with *t*, also the resting spring length is increasing linearly until age_*i*_ (*t*) = *t*_*g*_. Further, *t*_*d*_ (*i, t*) < *t* _*a*_, if cell *i* is undergoing apoptosis with *t*_*d*_ (*i, t*) → 0 with increasing time *t*. Thus, also *s*_*i*_ (*t*) → 0 and in total, 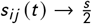.

#### Definition 5 (Linear Spring Force)

i. *The* force *exerted by cell i on cell j at time t is defined by the vector*

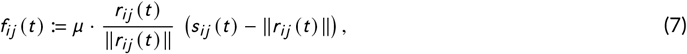

*where µ* ∈ ℝ_≥ 0_ *is the so-called* elasticity constant.
ii. *We further define the sum of all forces exerted on cell i at time t as*

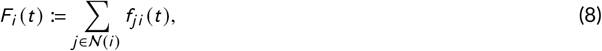

*where* 𝒩 (*i*) *denotes the Delaunay neighborhood of cell i, corresponding to the dual graph of the Voronoi tessellation of cell i*.

If *s*_*ij*_ (*t*) > ‖*r*_*ij*_ (*t*) ‖ in Equation (7), i.e., if the cells have not reached their natural distance, the force is called *repulsive*, driving the cells further away from each other. If *s*_*ij*_ (*t*) < ‖ *r*_*ij*_ (*t*) ‖, the force is called *attractive*.

Further, the parameter *µ* describes the elasticity of the cell. Large values of *µ* lead to greater forces, in other words, the cells are easier to push away or pull nearer, respectively. One might make this parameter dependent on the mutation state of a cell or on the cell type, which is due to future work.

In general, *f*_*ij*_ = − *f*_*j i*_, unless both cells are of different ages less than *t*_*g*_, i.e., both are newly divided cells from distinct mother cells. The force can now be used to define the motion of cells over time, where we assume Brownian dynamics, as in the model by ^13^. However, we extend the originally proposed equation of motion in ^13^ by a mechanism causing upward migration. This can be achieved by incorporating a basement membrane flow, additionally to mitotic pressure ^35^. The former is represented by an additive term increasing the second component of the cell position vector *r*_*i*_, as defined in the following.

#### Definition 6 (Equation of Motion)

*The change of the position r*_*i*_ *of cell i over time t is described by the ordinary differential equation*

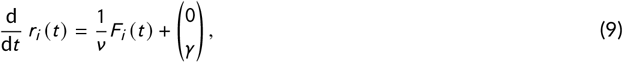

*where ν* ∈ ℝ_>0_ *is the so-called* damping constant *of a cell and F*_*i*_ (*t*) *are the forces exerted on cell i at time t defined in Definition 5. Further, γ* ∈ ℝ _≥ 0_ *is the additional increase in height caused by the basement membrane flow*.

The parameter *ν* describes cell-matrix adhesion. The greater the value of *ν*, the stronger the cell adheres to the extracellular matrix, which is in our case the basement membrane. This leads to decreased cell mobility. The latter is described by 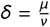 with *µ* defined in Definition 5.

In our case, the Equation of Motion (9) is discretized in time using the forward Euler method

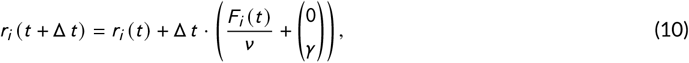

where Δ *t* denotes the default time step. This cell migration update is called at every time step, before checking for cell division.

### 2.5 | Modeling Cell Death

We will incorporate three different mechanisms how a cell of a crypt can die, where in each case, the respective node is removed from the Voronoi tessellation and after this process, all associated springs are deleted. The incorporated mechanisms are: 1) The cell is sloughed into the colonic lumen at the top of the crypt, 2) the cell falls victim to homeostatic mechanisms, or 3) the cell dies due to the acquisition of a disadvantageous mutation.

#### Modeling Cell Sloughing

The cells regularly undergo apoptosis after they finished the migration through the crypt. This is implemented in the model by letting all cells die when they reach the height of the crypt, i.e., if

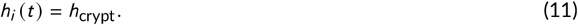

However, we currently do not model the survival of *APC* inactivated cells, which can lead to the formation of polyps. This could be circumvented by assuming a cell dependency, in particular a mutation state dependency, of this mechanism.

#### Modeling Homeostasis

We assume that homeostatic mechanisms may lead to cell death in order to avoid an overpopulation within the TA cell compartment. For this, we assume that cells with a surface area below a certain threshold *τ*_size_ are not viable and will die. Cells which have acquired either two *CTNNB1* mutations or two *APC* mutations are assigned a lower threshold, since these cells are partially able to ignore biochemical signals which would normally induce apoptosis. In formulas, this reads

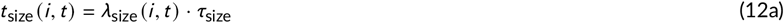

where

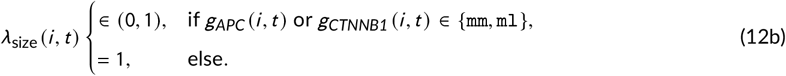

This drastically increases the chance of survival and eases proliferation for those cells.

We assume further homeostatic mechanisms within smaller subpopulations, like Wnt-induced senescence, and assign an additional probability *p*_*cc*_ of homeostatic death to all cells in the following way.

##### Proposition 1

*Let p*_*cc*_ ∈ [0, 1] *denote the probability of homeostatic cell death per cell cycle. Further, let* Δ*t denote the default time step and* 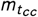 *the average duration of the cell cycle, both in hours. Then, the probability of cell death at each time step is given by*

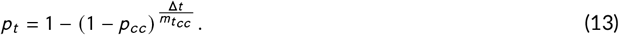

**Proof** The number of time steps per hour is given by 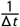. Therefore, there are 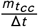 time steps per cell cycle. The probability of not dying during one cell cycle duration is

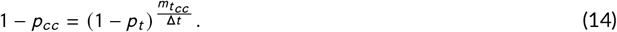

Rearranging for *p*_*t*_ yields the statement.

#### Modeling Mutation-Induced Death

Mutations damaging a cell in such a way that it is not viable anymore are called *lethal* mutations. We incorporate this mutation-induced death as apoptosis of the daughter cell. The probability of a lethal point mutation, *p*_mutdeath_, is higher in MMR-deficient cells, as those have a higher mutation rate. In formulas, we obtain

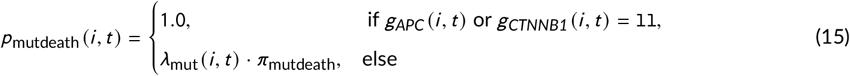

with *λ*_mut_ defined in Equation (4b) and the value of *π*_mutdeath_ used in the simulations given in Appendix B. Further, the effects of the different mutations on cell death are summarized in Table 1.

### 2.6 | Modeling Stem Cell Dynamics

We assume that each colonic crypt is populated by a single stem cell at a time, which populates the crypt for the time *t* _stem_ of stem cell cycle length. This active stem cell divides asymmetrically, renewing itself and giving rise to a new TA cell. If a mutation occurs upon cell division, the TA cell passes on the mutation to its descendants. In our model implementation, all cells in the current bottom row of the Voronoi tessellation become mutated, if the active stem cell becomes mutated. Adjacent to it, there are *S* − 1 ∈ ℕ quiescent stem cells, which do not divide. Over time, the *S* stem cells alternate at populating the crypt with a probability *p*_change_ of stem cell exchange per stem cell division. The new active stem cell is chosen with uniform probability among the *S* − 1 quiescent stem cells. Stem cell death is only possible due to a lethal mutation with a probability given in Equation (15). In this case, symmetric division of an adjacent stem cell compensates the dead stem cell, in order to maintain a constant number *S* of stem cells over time.

## 3 | SIMULATION RESULTS

All model simulations and *in silico* experiments are conducted within the Chaste framework (Version 2019.1) ^36^, which can be accessed from www.cs.ox.ac.uk/chaste. We adapted and extended the existing Chaste implementation with main changes in the cell cycle model and additional features for the LS-related mutations, for stem cell dynamics, and for the feedback mechanisms affecting homeostatic cell death. In particular, we added the three mutations of interest, and included the effects of each on the cell cycle and cell differentiation models. Further, we included our own stem cell model and extended the already existing cell migration model, as described in Section 2.4. Finally, the feedback mechanism was implemented in Chaste. In summary, our model is built on the general crypt model by Chaste, but some features were implemented from the ground up. The implementation is accessible on GitHub https://github.com/Mathematics-in-Oncology/ComputationalColonicCrypts/releases/tag/v1.0, release v1.0. A pseudocode of the computational model is given in the Appendix C.

The simulations were run on a modern workstation. As parallelization is not yet available in Chaste, each crypt was computed in a sequential way. For simulating a single crypt with 1600 cells for one year of human life-time based on our parameter setting, approximately 630 million operations had to be computed. This took approximately 44 hours of computational time.

For the simulation results, we analyze epithelial renewal times in mice and humans, monoclonal conversion of different types of mutations, as well as the influence of cell location and stem cell dynamics on the spread of mutations within a crypt. By this, we are able to gain a better understanding of the biomedical mechanisms leading to advantageous mutations within a crypt, which in turn are known as driver events in colorectal carcinogenesis. Further, we obtain *in silico* estimates for the duration of epithelial renewal and of monoclonal conversion of different mutations in these driver genes.

### 3.1 | Feedback Mechanisms and Epithelial Renewal in Non-Mutated Crypts

**Feedback mechanisms within the crypt** are modeled by homeostatic cell death in order to avoid an overpopulation of the crypt with TA cells. Here, only the parameters for modeling cell mobility and TA cell cycle length determine the size of the effect, which demonstrates the importance of feedback mechanisms for tissue organization and homeostasis. The feedback mechanisms are incorporated in our model by assuming exemplarily that cells smaller than a certain threshold are non-viable. The exact value of this threshold regulates the number of cells present upon the crypt’s homeostatic equilibrium. In other words, the minimum cell size determines the number of cells present in crypt homeostasis.

**The renewal time of a crypt** is the duration of the complete exchange of all cells within this crypt, which is rather short in many tissues. While in the murine small intestine, reliable estimates of less than one week have been established ^37^, estimations for the human colon are less precise.

In our model, the renewal time is most significantly influenced by the crypt size and the parameters describing cell mobility, in particular by the interplay of the overall cell mobility parameter *δ* and the basement membrane flow parameter *γ* introduced in Definition 6.

Using the parameter combination described in Appendix B, we estimate the renewal time of crypts *in silico*: A crypt which initially consists of about 200 cells, resulting in an equilibrium state of about 220 cells, is renewed every 6 days. This crypt size is representative of a murine crypt and the predicted renewal time is in concordance with the available estimates in mice ^38^. For a human colonic crypt, here initially consisting of 1600 cells, the *in silico* estimates for the renewal time increase to three weeks. This suggests that the process of epithelial renewal takes a number of weeks in humans. The estimates are illustrated in Figure 2.

**FIGURE 2.**
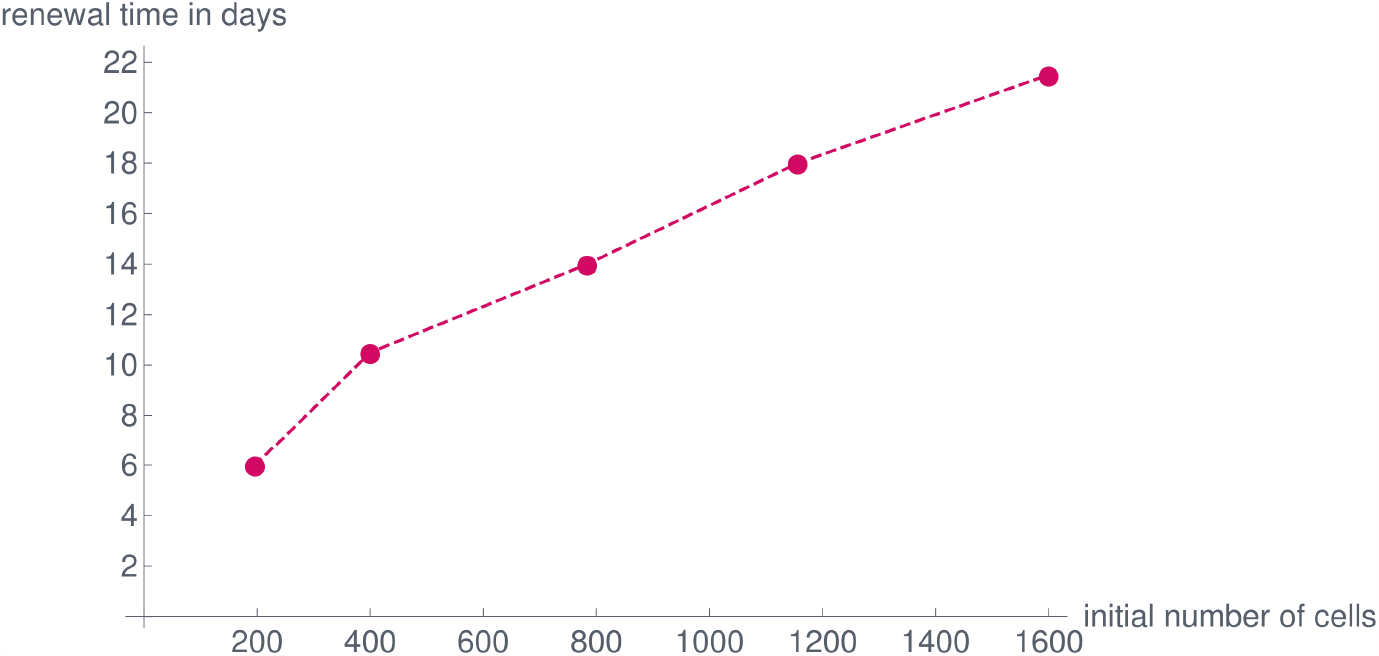
Renewal times depending on the initial cell number. Single simulations for an initial number of 196, 400, 784, 1156 and 1600 cells, respectively, result in equilibrium cell numbers of about 220, 460, 910, 1350 and 1850 cells. The initial setup in all cases followed a 4:1 ratio of crypt height to crypt circumference, measured by the number of cells. All other parameters were set as in Appendix B. All performed simulations showed renewal times only differing by a few hours.

### 3.2 | Influence of Stem Cell and TA Cell Mutations on Intra-Crypt Dynamics

We are able to analyze the effect of different types of mutational processes, namely non-transforming and transforming mutations. Exemplarily, we consider the following non-transforming mutations

⊳ two inactivating MMR mutations: *g*_MMR_ (*i, t*) ∈ {`mm, ml, ll`},
⊳ one inactivating *APC* mutation: *g*_*APC*_ (*i, t*) ∈ {`m, l`},
⊳ one activating *CTNNB1* mutation: *g*_*CTNNB1*_ (*i, t*) = `m`.

As examples of transforming mutations, we consider the double-hit *APC* or the double-hit *CTNNB1* mutation, i.e.,

⊳ *g*_*APC*_ (*i, t*) ∈ {`mm, ml`} or
⊳ *g*_*CTNNB1*_ (*i, t*) ∈ {`mm, ml`}.

#### 3.2.1 | Spread of Non-Transforming Mutations

For analyzing the spread and monoclonal conversion of non-transforming mutations, we initialize the computational model with the corresponding mutation in the active stem cell and run several simulations.

##### An MMR-deficient stem cell almost always leads to monoclonal conversion between 6 and 9 weeks

For analyzing the spread of MMR deficiency throughout the crypt, we use an MMR-deficient stem cell as initial condition and run 98 simulations. 88 of them predicted monoclonal conversion to be completed after between 42 and 71 days, with an average value of 51.9 days within this subset of the simulations, illustrated in Figure 3. In four of the remaining simulations, a biallelic *APC* mutation occurred before monoclonal conversion was completed. The predictions of the other six remaining simulations are discussed in Section 3.3.

**FIGURE 3.**
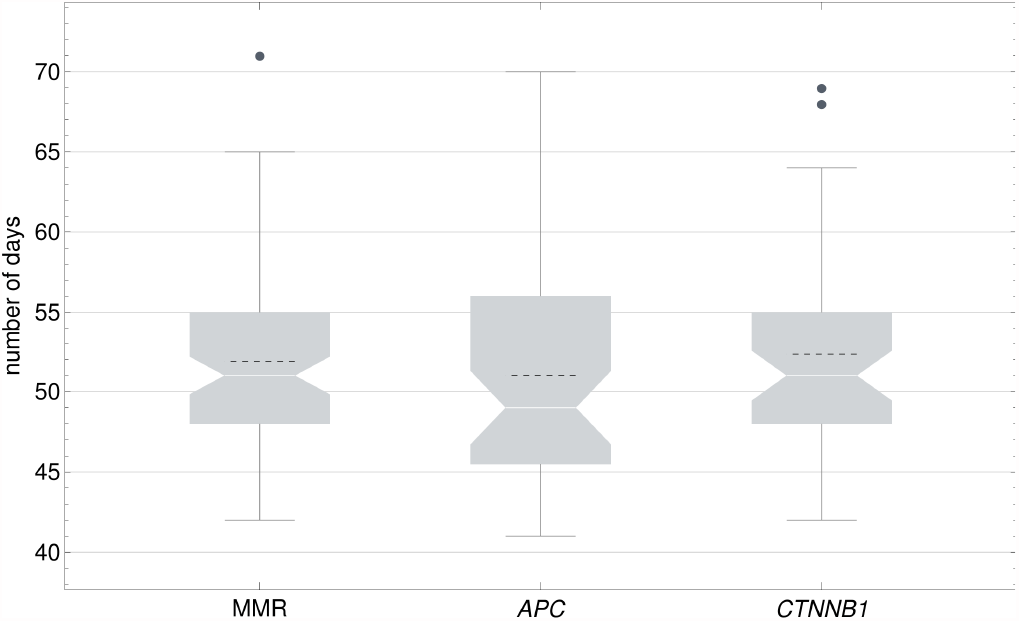
Comparison of monoclonal conversion of MMR deficiency, monoallelic *APC* and *CTNNB1* mutations. The results are illustrated by notched Box-Whisker plots, where the notches show the 95% confidence intervals and the dashed lines indicate the means. The respective medians (white line) amounted to 51 days (95% CI: [49.8; 52.2]) for MMR deficiency, 51 days (95% CI: [49.4; 52.6]) for monoallelic *CTNNB1* mutations, and 49 days (95% CI: [46.7; 51.3]) for monoallelic *APC* mutations.

##### The kinetics of the spread of monoallelic *APC* and *CTNNB1* mutations resemble the one of biallelic MMR mutations

After a stem cell mutation in either *APC* or *CTNNB1*, the mutation spread throughout the crypt in the great majority of cases (*APC*: 52/53, *CTNNB1*: 50/54). For both types of mutations, our model predicts monoclonal conversion to take about 52 days on average (*APC*: 51 days, *CTNNB1*: 52.3 days), see Figure 3.

Based on a Kruskal-Wallis test, the null hypothesis that the means for MMR, *APC* and *CTNNB1* are the same is not rejected at the 5% significance level (*p* = 0.266) indicating a highly similar duration time of monoclonal conversion for non-transforming mutations. The similarity resides in the fact that the delay of differentiation of cells harboring the monoallelic *APC* or *CTNNB1* mutation does not provide any advantage regarding the *speed* of the spread of the mutation, as it does not change the cell’s proliferative behavior, survival, or mobility. This also implies a high similarity of the duration time of monoclonal conversion for non-transforming mutations and wild-type crypts. The effect on differentiation only increases the probability that a mutation spreads at all. This probability is indeed very high for stem cell mutations, but rather low for TA cell mutations. Here, most of the expansions of a mutated clone can be prevented by the feedback mechanism described above, and by the fast renewal of the crypt. The latter process results in such clones being frequently washed out of the crypt, which becomes inevitable as soon as the cells of the clone complete differentiation. This underlines the importance of stem cells regarding the origin of colorectal cancer. **Monoallelic *APC* mutations in TA cells** are frequent in our simulations due to the high number of hotspots. Hotspots are regions of a gene which give rise to a phenotypical change if mutated (see Appendix B). As we assume no dominant-negative effects of *APC* mutations for modeling, monoallelic *APC* mutations are rather non-transforming since the second allele continues to produce a sufficient amount of the APC protein. In addition, differentiation is prolonged due to the lower Wnt threshold. The expansion of a monoallelically mutated *APC* clone can be prevented in most cases by the feedback mechanisms and the fast renewal of the crypt resulting in a wash-out.

In the presence of a biallelic MMR mutation, we assume point mutations to be much more likely than in normal cells. In the case of an MMR-deficient stem cell, this results in the formation of many clones with monoallelic *APC* mutations, most of which are however washed out. For instance, we observe up to 45 monoallelic *APC* mutations in a single crypt over the course of 30 weeks, none of which gives rise to a second inactivation event.

#### 3.2.2 | Spread of Transforming Mutations

If a cell with one mutation in either *APC* or *CTNNB1* acquires a second hit, it becomes partially resistant to apoptosis. Further, the minimum size threshold for homeostatic death is lowered, which both result in a heavily increased chance of survival. The latter is reflected by the spread of such mutations.

##### Biallelic *APC* and *CTNNB1* mutations always lead to monoclonal conversion

The evasion of the feedback mechanism and of differentiation allow the clone with biallelic *APC* or *CTNNB1* mutations to initially grow exponentially and always take over the crypt in our simulations. Exemplary figures and simulations illustrating the evolution of cells with biallelic *APC* or *CTNNB1* mutations are shown in Figure 4 and in Appendix D. As an important contrast to non-transforming mutations, the location of the first mutated cell does not play a role regarding the mutation’s ability to spread. Further, monoclonal conversion is completed significantly faster compared to the non-transforming mutations: The means are 21 days and 21.6 days for biallelic *APC* and *CTNNB1* mutations, respectively, see Figure 5. However, based on a Kruskal-Wallis test, there is no significant difference in the means between the biallelic *APC* and *CTNNB1* mutations (*p* = 0.09).

**FIGURE 4.**
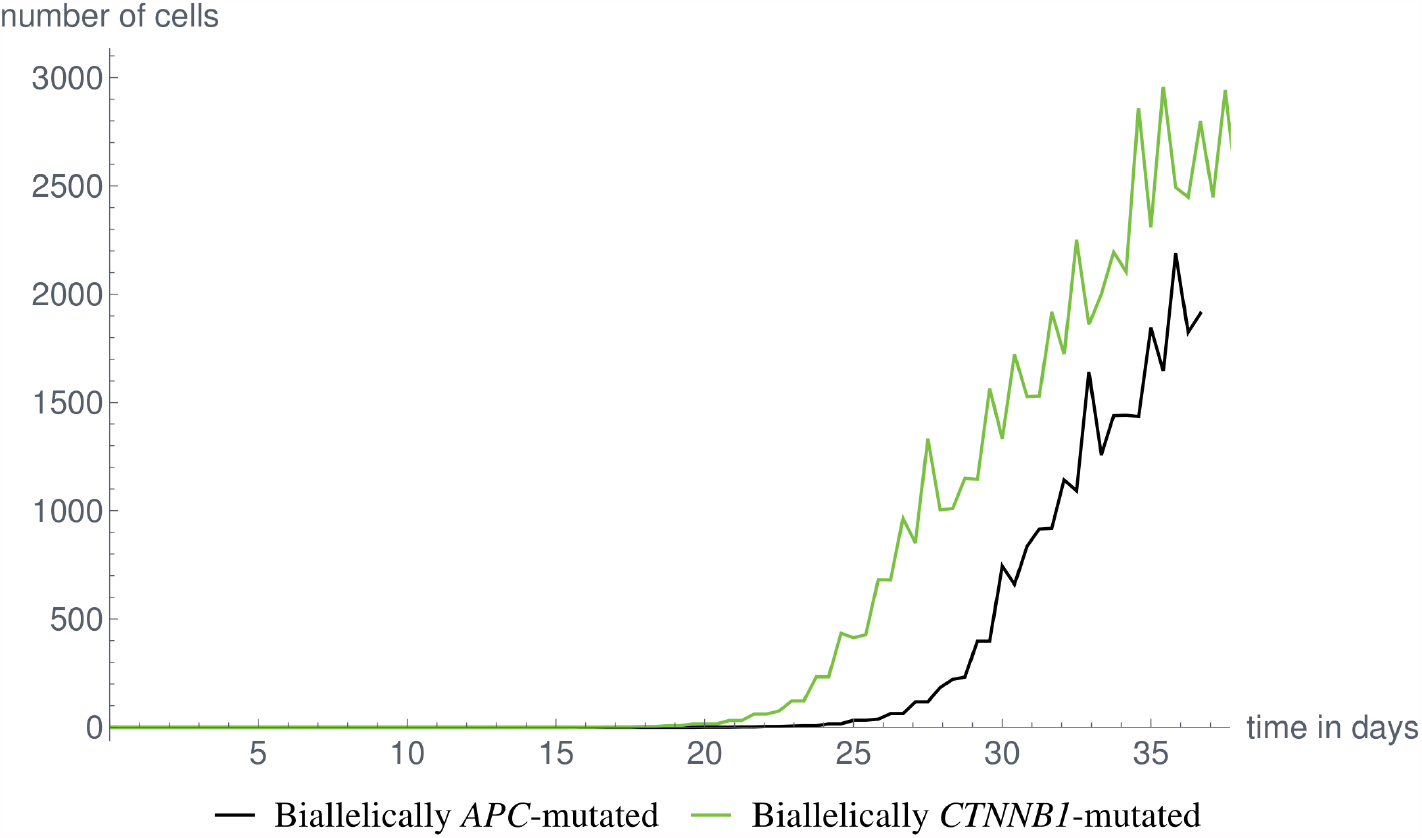
Exponential expansion of a biallelically mutated clone. Evolution of the number of biallelically *APC*-mutated cells (black). and of biallelically *CTNNB1*-mutated cells (green). While the simulation for *CTNNB1* ran for approximately 60 days and it reached the plateau after 35 days, we only show the initial 35 days to ensure a one-to-one comparison. Videos of the complete simulations are provided in Appendix D.

**FIGURE 5.**
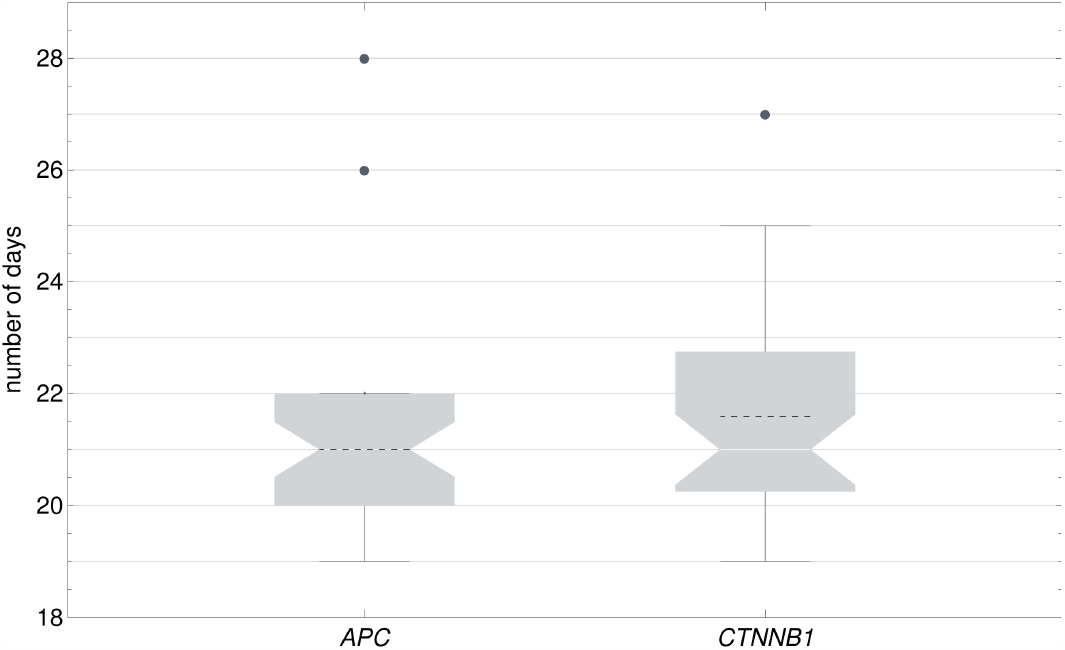
Comparison of monoclonal conversion of biallelic *APC* and *CTNNB1* mutations. The results are illustrated by notched Box-Whisker plots, as in Figure 3. For both biallelic *APC* and *CTNNB1* mutations, the median over our simulations (41 for *APC*, 39 for *CTNNB1*) amounted to 21 days with 95% confidence intervals of [20.5; 21.5] and [20.4; 21.6], respectively.

It might be feasible to assume that each monoclonal crypt will give rise to an aberrant lesion such as an adenomatous polyp. Importantly, this would directly imply that any biallelic *APC* or *CTNNB1* mutation occurring in a cell of the crypt results in the development of such a lesion.

### 3.3 | The Influence of Cell Location on Mutation Spread

The spread of a mutation is predicted to depend on the location of the cell, in which the mutation first occurs ^39^. Within our short-term simulations, a clear trend in favor of the crypt base could be observed, which will be analyzed in the following sections.

#### The crypt base is the most stable environment within the crypt for mutated and wild-type cells

Mutated clones which originate in the lowest region of the crypt usually have more time to expand without facing complete differentiation. Consequently, the ability that mutations, especially non-transforming ones, manage to spread throughout the crypt at all increases when decreasing the position of the initial mutated cell. In our *in silico* experiments, we observe this for stem cell mutations, which are located at the bottom of the crypt and in most cases become monoclonal. As an additional example, a cell which acquires a monoallelic *APC* mutation near the bottom of the crypt consistently gives rise to larger clones, compared to cells located higher in the crypt. Examples are illustrated in Figure 6 and in Appendix D.

**FIGURE 6.**
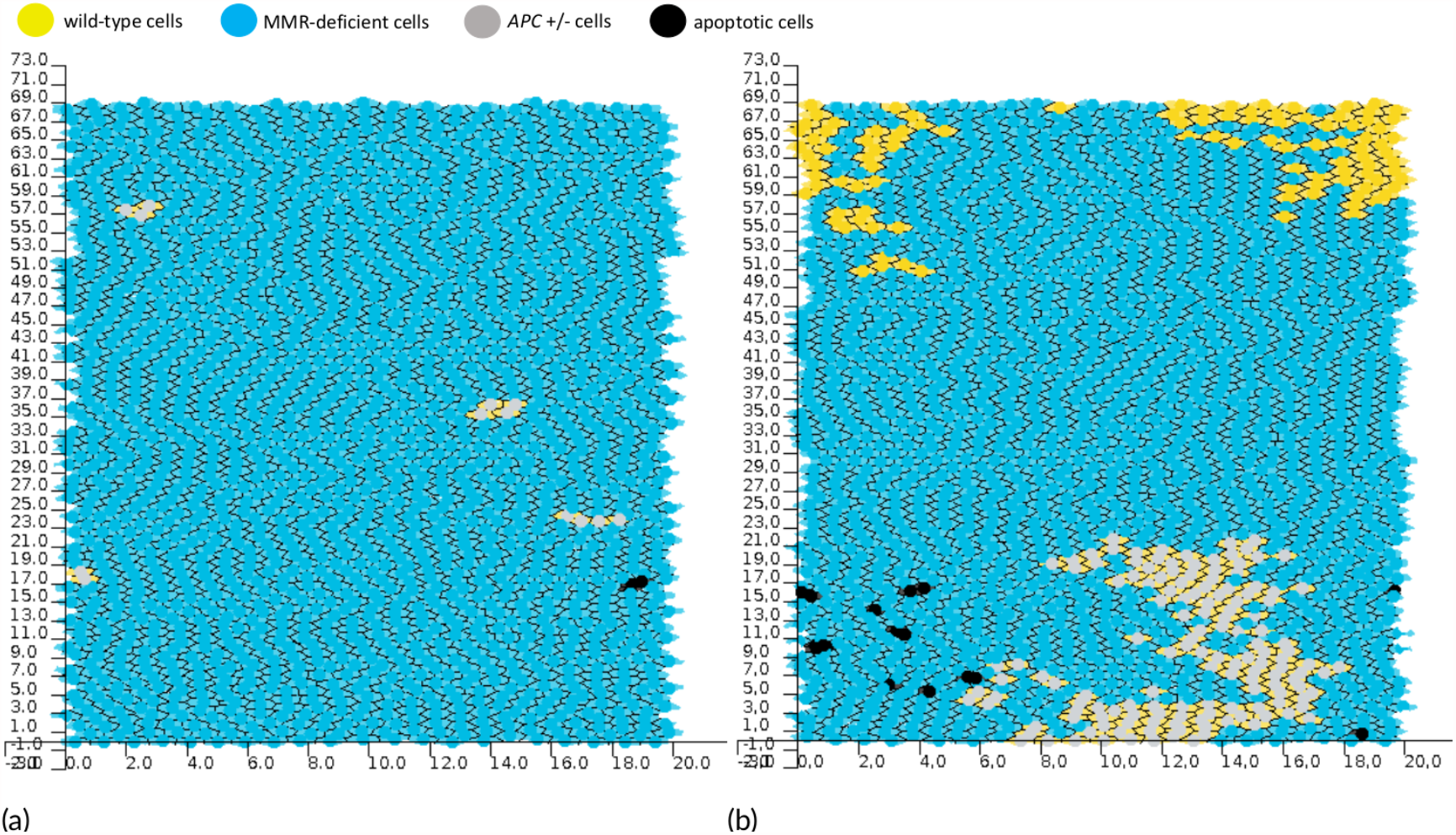
Spread of *APC* mutations dependent on cell position. Video frames of two separate simulations are shown, both crypts are already MMR-deficient by initialization. In all simulations, MMR-deficient cells are shown in blue, MMR-proficient cells in yellow, monoallelically *APC*-mutated in light gray, biallelic *APC*-mutated in dark gray with a black nucleus, biallelically *CTNNB1*-mutated in green and apoptotic cells in black. The higher the *APC* mutations occur, the smaller are the resulting clones. (a) Four small monoallelically *APC*-mutated clones are visible 64 days after the start of the simulation. All mutations occurred in the lower middle part of the crypt. (b) A monoallelic *APC* mutation occurred after 20 days in the lowest cell row of TA cells. This led to the formation and temporal persistence of a large monoallelic *APC*-mutated clone, here shown after 40 days. Videos of the complete simulations are provided in Appendix D.

However, the property of the crypt base being a stable environment is not limited to mutated cells. It can equally serve as a beneficial region for wild-type cells. In our simulations, a mutated stem cell only rarely does not give rise to monoclonal conversion. In this rare case, a clone of wild-type cells is able to persist for a sufficiently long period of time at the crypt base. This slows down the spread of the mutation, such that monoclonal conversion is either completed much later than on average, or never completed at all, explaining the outliers in Figures 3 and 5. An example of the former case is illustrated in Figure 7. We observed those delays of monoclonal conversion at least once for monoallelic stem cell mutations in both *APC* and *CTNNB1*, as well as in an MMR-deficient stem cell. If monoclonal conversion of these mutations is prolonged until the end of the cell cycle of the stem cell which populates the crypt after a stem cell mutation, the process might not be completed due to an exchange of the stem cell which populates the crypt.

**FIGURE 7.**
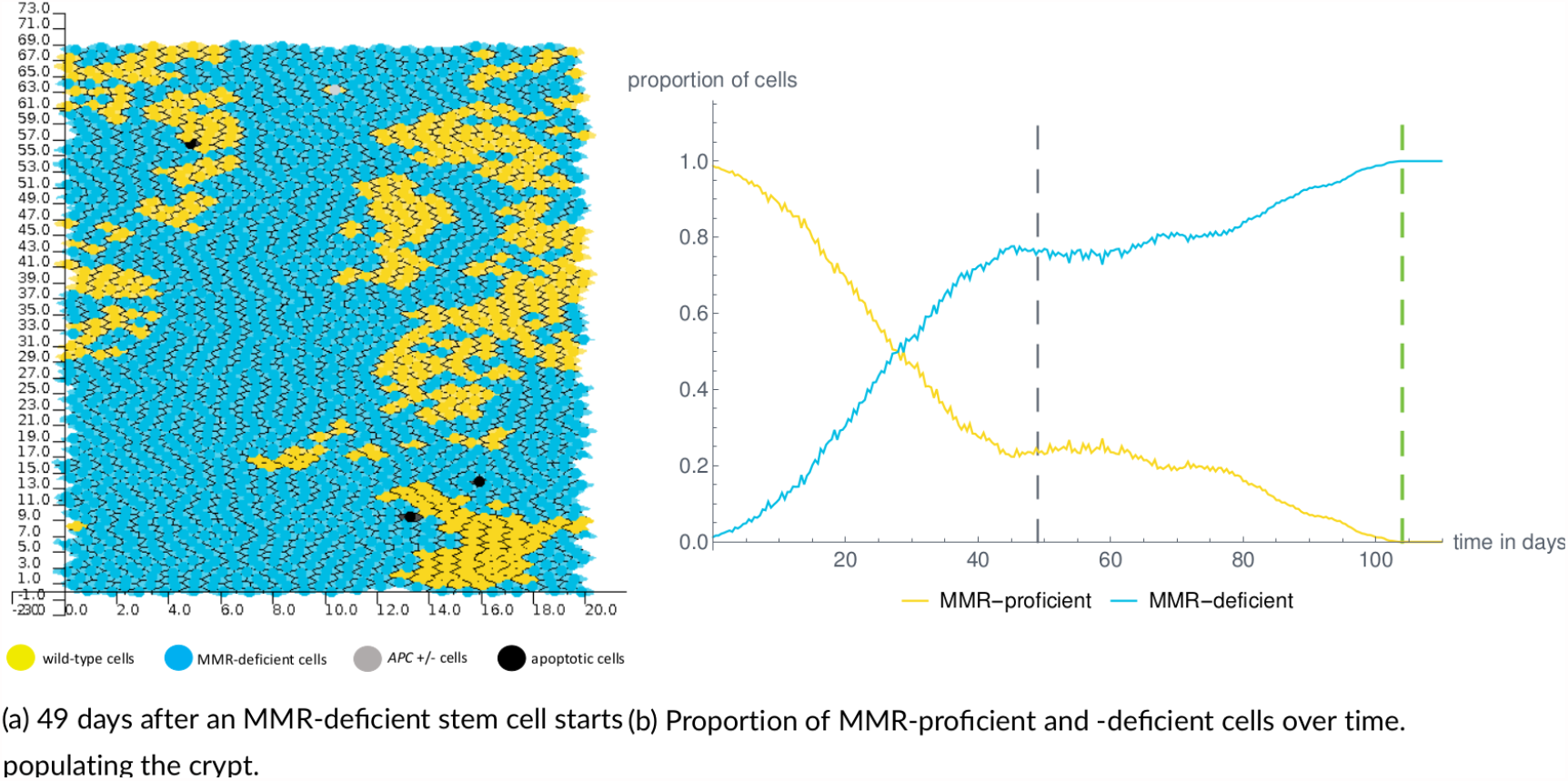
Prolonged monoclonal conversion of an MMR stem cell mutation. Color legend and initial condition are as before. A video of the complete simulation is provided in Appendix D. (a) After 49 days, close to the average duration of monoclonal conversion of MMR deficiency (see Figure 3), a large clone of MMR-proficient TA cells still resides near the crypt base. Note that many other differentiated MMR-proficient cells also have not been washed out at this point in time. As illustrated in Appendix D, three weeks later, the progeny of the same clone has lost the position near the crypt base and is subsequently washed out of the crypt. Monoclonal conversion was completed after 104 days (dashed green line). (b) The evolution of both cell types over time. Note that the duration of monoclonal conversion is substantially longer compared to the average in Figure 3.

#### Top-down vs bottom-up morphogenesis is determined by the Wnt threshold

During our simulations, we observed two ways of morphogenesis of biallelically *APC*-mutated cells: 1) The second hit occurs in parts of the crypt where wild-type cells already have completed differentiation. In this case, as a result of the direction of cell migration, the mutation spreads toward the top of the crypt before taking over the bottom half, see Figure 8 (a), (b) and Appendix D. This is consistent with *top-down morphogenesis*, as discussed in Appendix A. 2) Due to the proliferative zone of the crypt being located in its lowest quarter, the majority of biallelic *APC* mutations are predicted to originate in the bottom region. This is known as *bottom-up morphogenesis*, illustrated in Figure 8 (c), (d) and Appendix D. In our simulations, the spread of a biallelically *APC*-mutated clone follows top-down morphogenesis in 22% of cases (9/41) and bottom-up morphogenesis in 66% of cases (27/41), while the remaining five cases could not be clearly identified as one or the other. Altogether, our *in silico* experiments show both modes of morphogenesis to be possible.

**FIGURE 8.**
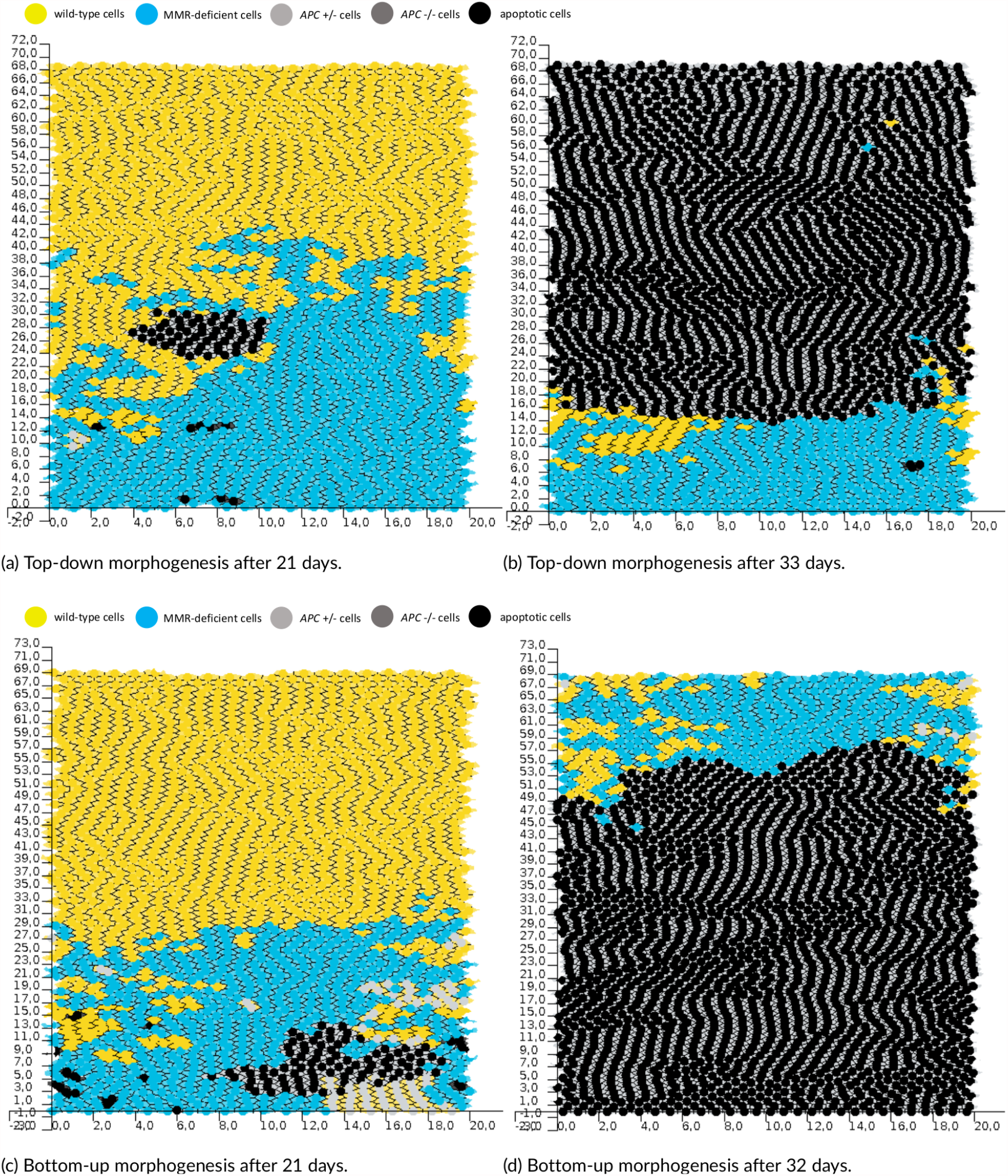
Different modes of morphogenesis of a biallelically *APC*-mutated crypt. Color legend and initial condition are as before. (a), (b): The second *APC* mutation occurred in the region of wild-type FD cells after 15 days. The biallelically *APC*-mutated clone consists of 56 cells after 21 days (a) and of about 1800 cells 12 days later (b). The upper parts of the crypt are populated first, before the bottom quarter. The second *APC* mutation occurred after 14.5 days in a cell within a monoallelically *APC*-mutated clone consisting of 24 cells, which can be seen below the biallelically *APC*-mutated clone. The latter consists of 90 cells after 21 days (c) and of about 1900 cells 11 days later (d). Videos of the complete simulations are provided in Appendix D.

Based on our analyses, the proportion of biallelically *APC*-mutated crypts, which develop in a top-down manner, depends most notably on the Wnt threshold *τ*_wnt_ for monoallelically *APC*-mutated cells. The lower this value, the later in the process of migration these cells differentiate and thus the higher the frequency of top-down monoclonal conversions. However, lowering the Wnt threshold also gives rise to more biallelically *APC*-mutated crypts overall.

### 3.4 | The Effect of Stem Cell Exchange on Monoclonality

The effect of a stem cell exchange event on the monoclonality of a crypt depends on the respective mutation.

#### Stem cell exchanges can lead to transient polyclonality

Our model simulations suggest that monoclonal biallelic MMR mutations are regularly washed out of the crypt after a stem cell exchange event. This process is illustrated in Figure 9. The observations are in concordance with our biological understanding, since MMR deficiency per se is not expected to provide a proliferative advantage to the cells and the transition to an alternative monoclonal status with transient polyclonality is likely to appear.

**FIGURE 9.**
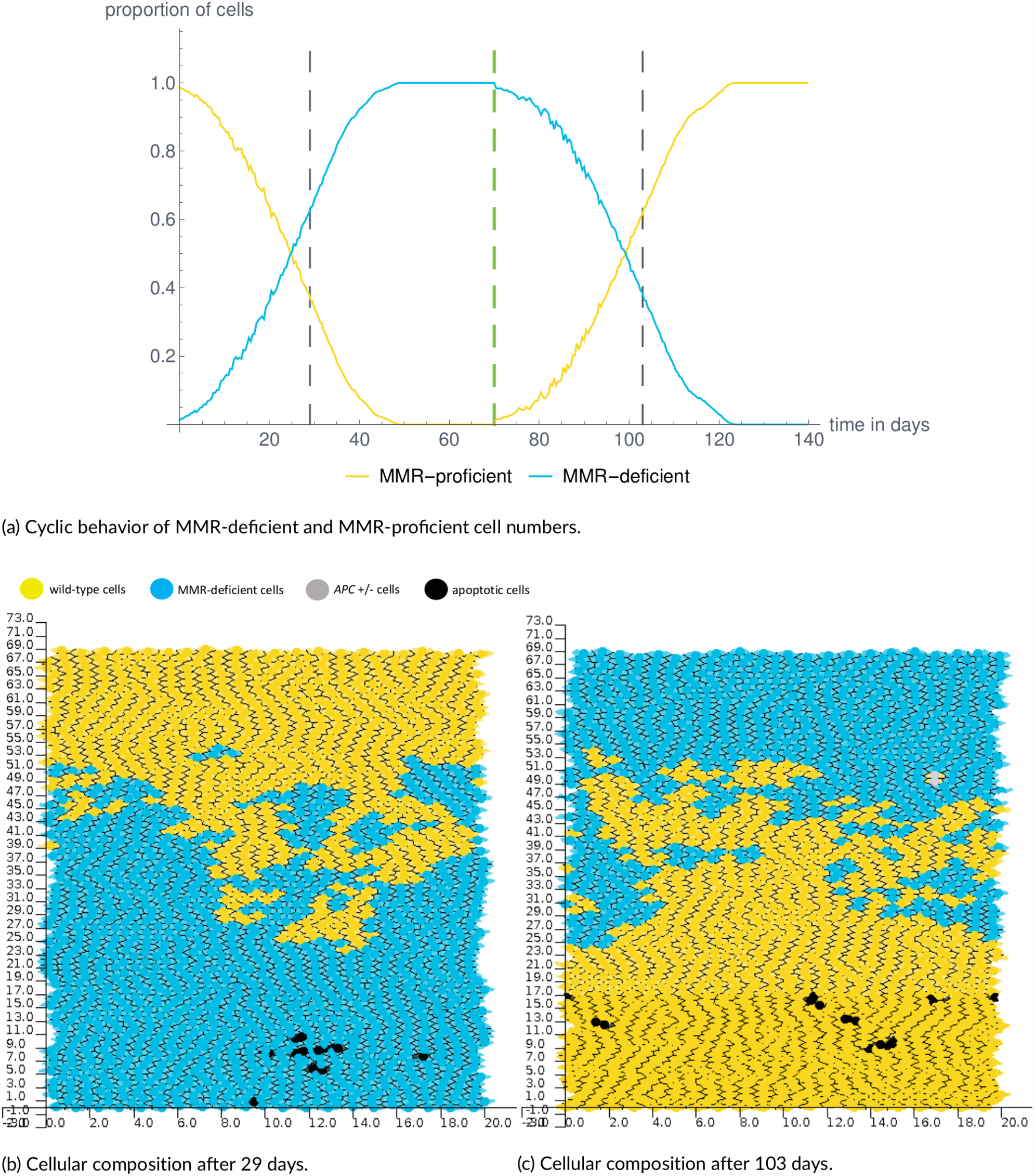
Development and loss of MMR deficiency due to stem cell exchange. (a) Initialization with an MMR-deficient stem cell leads to monoclonal conversion of MMR-deficient cells (blue curve) after 49 days, while the wild-type cells (yellow curve) are washed out of the crypt. After a first stem cell cycle of 70 days (green line), a wild-type stem cell populates the crypt and the process of monoclonal conversion is reversed. After an additional 54 days, no MMR-deficient cells are left in the crypt. (b) and (c) show the cellular composition of the crypt after 29 days and 103 days, respectively. The time points are illustrated in (a) by gray lines with colors as before. A video of the complete simulation is provided in the Appendix D.

For monoclonal monoallelically *APC*-mutated or *CTNNB1*-mutated crypts, we observe similar scenarios. The underlying reason is again that the delay of differentiation caused by these mutations does not yield a significant advantage compared to the expansion of MMR-deficient or wild-type cells.

#### Monoclonality can persist upon stem cell exchanges

This is true for the monoclonality of biallelically *APC*-mutated or *CTNNB1*-mutated crypts. In our *in silico* experiments, the division of a wild-type stem cell populating the crypt after a stem cell exchange gives rise to a small population of wild-type TA cells. However, this population becomes extinct after 2–3 days. Since the biallelic mutations provide a significant proliferative advantage, wild-type cells are inferior to such cells and do not have the ability to recapture the crypt. Together with our observations from the previous sections, this implies that biallelic *APC* or *CTNNB1* mutations in any cell within the crypt result in the development of a monoclonal crypt, and that this crypt persists over time.

## 4 | DISCUSSION

We presented a computational model describing the evolution of colonic crypts in LS scenario. It uses a cylindrical representation of the crypt with a Voronoi tessellation for the cells. Based on the existing models for cell differentiation and cell migration, we developed a novel approach for stem cell dynamics and for cell death which occurs due to feedback mechanisms. For the former, we assume one of the current biological hypotheses that the whole crypt is populated by a single stem cell at a time ^16;17^ which coexists with quiescent stem cells at the crypt bottom.

The use of a Voronoi tessellation model was shown to be well-suited to model epithelia ^29^ for several reasons. Voronoi tessellation models are suited for both short-range and long-range dynamics due to their smooth transitions and the ability to easily verify cell neighborhoods using its dual graph, the Delaunay triangulation.

Our computational model provides the ability to investigate a variety of biomedical aspects of colonic crypt evolution which are crucial for its detailed understanding. Besides studying the evolution of non-mutated crypts, it gives insights into the evolution of crypts carrying possible driver mutations and initiating events of LS carcinogenesis. In particular, we analyzed the effect of biallelic *CTNNB1* mutations, which are recently demonstrated ^18^ to be driver events particularly in LS carcinogenesis. We analyzed the spread and monoclonal conversion of different driver mutations and the effect of stem cell dynamics thereof. All of these non-mutated and mutated crypts were examined by varying initial conditions, such as somatic stem cell mutations or pathogenic germline variants in different MMR genes. By a suitable choice of parameters and further programming, the model could also be used to simulate the development of other types of colorectal cancer. By initializing the crypts with no pathogenic germline variants, sporadic CRC and Lynch-like carcinogenesis ^19^ could be modeled. By initializing with a pathogenic germline variant in *APC*, the model could be used to model CRC initiation in FAP. The presented technique of incorporating point mutations and LOH events into cell-based models could be used not only in other CRCs but also for cancer in other organs. The mutation and LOH model based on gene-dependent mutation rates and mutational events which are independent of or dependent on other mutations was already adapted exemplarily for FAP and sporadic CRC in ^32^, where it is used for modeling the evolution of the number of crypts.

Concerning the dynamics of non-mutated tissue, the simulation results predicted that feedback mechanisms are necessary to ensure crypt homeostasis. However, the specific mechanism is currently not known from a biological point of view and our implementation (i.e., the death of small cells) should be regarded as exemplary, although the loss of cell integrity due to overwhelming pressure appears to be reasonable. Once further biological hypotheses are available, they can be included in the model.

Furthermore, our estimates obtained for both human and murine crypts suggest that the process of crypt renewal might take several weeks in humans, where we identified the magnitude of cell mobility and the crypt size as the two determining parameters. While the estimates are in concordance with experiments in mice, the *in silico* duration in humans is yet to be experimentally validated. Further, the renewal times for different initial cell numbers are not strictly linear. This is in concordance with our expectation since 1) the crypt height, which is one determining factor, does not increase linearly with the total cell number and 2) the increase in the number of FD cells inhibits crypt renewal more than the same (relative) increase in the number of TA cells accelerates it.

The majority of our simulations analyzed the spread of different mutations within the crypt, either occurring in a stem cell or a TA cell. The main focus was on the duration of monoclonal conversion, distinguishing between different types of mutations: non-transforming and transforming mutations. Our model predicted the former to become fixated more slowly than the latter, with estimates ranging between 6 and 9 weeks, and 2 to 4 weeks, respectively. 88/98 simulations predicted monoclonal conversion of MMR deficiency initiated by an MMR-deficient stem cell to be completed after between 42 and 71 days, with a median of 51 days (95% CI: [49.8; 52.2]). The reported predictions of time span required for a monoclonal conversion of a crypt carrying certain mutations provide the basis for future studies, in particular addressing the time required for an aberrant crypt to become an endoscopically visible lesion.

Our estimates for non-transforming mutations differ from the mean value of 18.6 days reported in ^15^. However, in the latter study, the spread of a non-transforming mutation throughout a crypt consisting of 250 cells was examined with a significantly shorter average stem cell cycle duration of 24 hours compared to a few weeks in our approach. These factors may explain the difference between the estimates.

We further discussed the influence of stem cell dynamics on monoclonality. With our simulations, stem cell exchange can restore the integrity of crypts and contribute to the elimination of mutations without a directly transforming effect, including MMR gene mutations. This suggests that it serves as a mechanism which can inhibit carcinogenesis. Furthermore, the results of our simulations of MMR-deficient crypts may help explain why many MMR-deficient crypts do not progress to larger lesions: Such crypts might be detectable via staining, but frequently lose their status due to stem cell exchange later on. However, it is important to note that the monoclonality might be regained, in case the mutated stem cell populates the crypt again. This pattern of loss and recovery of monoclonality might occur repetitively, as long as no further advantageous mutations occur, and the mutated stem cell does not die. As in LS individuals, a cell only needs a second hit to be MMR-deficient, this finding is of particular importance for these people. MMR-deficient crypts are much more likely to occur but not all of them might evolve into a carcinoma later due to the stem cell exchange mechanisms.

The same applies for biallelic *APC* or *CTNNB1* mutations. We observed that they lead to a monoclonal crypt which can develop to a clinically manifest lesion. Individuals with FAP carry a monoallelic germline mutation in the *APC* gene. This means, any second *APC* inactivation event is predicted to potentially result in the formation of a polyp. This is in concordance with the hundreds to thousands of adenomatous polyps frequently detected in FAP patients ^40^. Besides that, the simulations showed both top-down and bottom-up morphogenesis to be possible for biallelically *APC*-mutated cells whereby bottom-up morphogenesis was more frequent (66% of the simulations). This finding is in concordance with ^31^.

In summary, our results provide first mathematical clues for effective surveillance protocols for LS carriers. First, the duration of monoclonal conversion for different driver events is a first hint for the duration of single steps in LS carcinogenesis. By long-term simulations, we will address the question when these events happen and by this, our estimates could contribute to the question of surveillance intervals. Second, the possibility of disappearing MMR-deficient crypts is currently a medical hypothesis linked to the question of overdiagnosis in LS ^41^. During our simulations, we observed this scenario supporting the medical hypothesis. Future work will include more simulations for analyzing this effect in more detail.

Although our approach takes into account many different processes and mechanisms occurring within colonic crypts, several aspects are neglected or not modeled in great detail. While we concentrated on the Wnt pathway with high evidence regarding its role in colorectal cancer, the implementation of other signal gradients like Eph/ephrin and Notch signaling is possible with the presented model framework and is due to future work. Further, as we focused on the initiating events of Lynch syndrome carcinogenesis, mutations considered to occur at more advanced stages and/or less frequently in LS carcinogenesis, such as *KRAS* or *TP53*, were not implemented in our model, although in principle our model would allow the implementation of additional mutational events.

We carefully evaluated the level of complexity with the computational costs of our model regarding computation time and storage. Future work shall add further modeling complexity where needed while trying to not increase the computational expense to allow for long-term simulations. For example, this could include a dynamic cell cycle model, a detailed model for feedback mechanisms and further, possibly yet unidentified mutations frequently occurring in colorectal carcinogenesis.

## Conflict of Interest

The authors have declared that no conflict of interest exists.

## Data Availability

All relevant data are within the manuscript and its Supporting Information files.

## Supporting Information

We provide supporting information in the appendix of this manuscript including detailed biological background information for the crypt modeling, a description of the parameter setting in the model based on existing biomedical data, a pseudocode of the presented computational model, as well as legends to videos of the simulations. The videos are provided by additional files. All model simulations and *in silico* experiments are conducted within the Chaste framework (Version 2019.1) ^36^, which can be accessed from www.cs.ox.ac.uk/chaste. We adapted and extended the existing Chaste implementation with main changes in the cell cycle model and additional features for the LS-related mutations, for stem cell dynamics, and for the feedback mechanisms affecting homeostatic cell death. The implementation is accessible on GitHub https://github.com/Mathematics-in-Oncology/ComputationalColonicCrypts/releases/tag/v1.0, release v1.0.

## APPENDIX

### A | BIOLOGICAL BACKGROUND

In order to build a model for the crypt dynamics, we first need to understand the biological structure and functioning of colonic crypts. Our main focus lies on their cellular composition and the signaling pathways regulating fundamental dynamics such as cell proliferation, migration and differentiation.

#### Colonic Crypts

Colonic crypts are tubular invaginations within the colonic epithelium which are believed to be the origin site of CRC ^5^. According to current estimations, human colonic crypts are about 75 to 110 cells long and have an average circumference of 23 cells ^6^. Naive multiplication therefore suggests the total cell number per crypt to range between 1.7 and 2.5 thousand.

The cells of the colonic crypt can be divided into three groups: *stem cells, transit-amplifying cells* and *fully differentiated cells*. They are characterized by their functions and abilities with respect to the cell proliferation, division, differentiation and migration. To understand the differences between the cell types, we first have to describe the main aspects of the processes, starting with the cell cycle crucial for cell division.

#### The Cell Cycle

In order to sustain the integrity of tissues, it is crucial that cells are able to divide and grow, which we collectively term *cell proliferation*. Between two divisions, each cell undergoes a series of events known as the *cell cycle*, which includes the preparation for DNA replication (*G1 phase*), DNA replication (*S phase*), the preparation for cell division (*G2 phase*), and cell division (*M phase*) ^42^. The success of the cell cycle depends on complex signaling cascades consisting of proteins, enzymes and hormones, each fulfilling different tasks. The inhibition of single elements of these cascades can lead to the arrest of the cell cycle, inhibiting proliferation. In this case, the cell is said to be *quiescent*, or in the *G0 phase*.

#### Cell Differentiation

Cell differentiation describes the transition from one cell type to another, involving changes in size, shape, and responsiveness to biochemical signals ^43^. In most cases, this leads to the cell becoming more specialized, i.e., it fulfills more specific tasks within its respective tissue. A division of a cell which results in two new cells of the same type is called *symmetric*, while a division resulting in the creation of one cell of the same type and one cell of a more differentiated cell type is called *asymmetric*. Biochemical signaling cascades heavily influence the process of cell differentiation. One of the primary regulators might be the so-called *Wnt pathway* ^20^.

#### The Wnt Pathway

The Wnt pathway describes a complex signaling cascade involved in several distinct processes across all animal species, such as embryonic development, tissue regeneration and carcinogenesis. The ongoing activity of the Wnt pathway can prolong differentiation ^5^. This is due to the role of the two proteins APC and *β* -catenin: Broadly speaking, APC is part of a complex of proteins which degrades *β* -catenin and thereby prevents it from traveling to the cell nucleus, where it can induce cell division. The activation of the Wnt pathway involves blocking APC from degrading *β* -catenin and leads to ongoing cell division. Due to this connection, mutations in the *APC* gene and in *CTNNB1*, the gene encoding for *β* -catenin, have been linked to various types of cancer ^44^

#### Cell Migration

Beside biochemical signaling cascades, the function of a cell is heavily influenced by the interaction with other cells and with the extracellular matrix, which are all macromolecules in the intercellular space. These two types of interactions are collectively termed *cell adhesion*. The attachment of cells to the extracellular matrix is necessary for the directed movement of cells, termed *cell migration*. In the colonic crypt, the adherence of cells to the extracellular matrix causes an upward migration. Further, the division of adjacent cells creates a so-called *mitotic pressure*, which also contributes to cell migration ^35^. The latter is essential for the maintenance of structures, and the formation and regeneration of tissues within organisms.

#### Mutations

Upon DNA replication, errors can occur and, if not corrected, manifest as *mutations*, of which there are two broad classes: So-called *point mutations* only affect a single nucleotide, while *loss of heterozygosity* (LOH) refers to the loss of some region in one copy of the diploid genome, which can result in the deletion of whole genes.

As we consider LS colorectal carcinogenesis, we focus on mutations in the MMR genes with a pathogenic germline variant. For modeling the Wnt pathway, we include *APC* and *CTNNB1* mutations in the model, where *APC* is a classical tumor-suppressor gene with two mutations necessary for inactivation. In particular, we ignore a possibly dominant-negative effect of *APC* mutations resulting in a single hit necessary for inactivation ^45^. Further, in CRC, biallelic mutations of *CTNNB1* seem to be required to mediate an oncogenic driver effect ^18;46^

We can now explain the three main cell types of colonic crypts with their characteristics.

#### Stem Cells

Stem cells reside at the crypt bottom ^6^. These cells are undifferentiated and have unlimited proliferative potential, such that they can renew themselves and give rise to more differentiated progeny. According to ^16;17^, we assume that there is one active stem cell at a time populating the crypt, whereby the others are quiescent. In general, the stem cell cycle is much longer than the one of TA cells. Usually, stem cell division is *asymmetric*, leading to one stem cell and one transit-amplifying cell. During this process, mutations can happen which can also lead to mutation-induced death of the stem cell. In this case, one of the adjacent stem cells divides symmetrically leading to two new stem cells and a fixed number of stem cells over time. The relative frequency of either mode of division is a current research topic ^47^

#### Transit-Amplifying (TA) Cells

Transit-amplifying (TA) cells are located above the stem cells in the lower and lower middle part of the colonic crypt. These cells are thought to possess certain properties of both stem cells and fully differentiated cells, depending on how far along they are on the path to differentiation ^7^. TA cells lack the ability to endlessly regenerate as they only divide a certain number of times before becoming fully differentiated ^8^. We ignore the possibility of de-differentiation of TA cells in the current modeling approach. However, TA cells divide more frequently than stem cells. The mode of division is, among others, determined by the activity of the Wnt pathway within these cells. According to ^28^, constitutive activation of Wnt in cells leads to an expansion of the populations of undifferentiated cells, and reduced Wnt signaling results in a complete loss of undifferentiated cells. Further, mutations can happen during each division leading to potential mutation-induced death. Besides that, we assume that TA cells can die due to high mitotic pressure.

#### Fully Differentiated (FD) Cells

Fully differentiated (FD) cells are located above the TA cells and never divide ^42^, thus no mutations are generated. Due to cell adhesion and mitotic pressure, they migrate to the top of the crypt, where they undergo apoptosis and are shed into the colonic lumen.

#### Morphogenesis of Mutated Colonic Crypts

Whenever a mutation occurs in a cell of a crypt, this cell has the potential to pass on the mutation to its progeny, possibly resulting in almost all cells in the crypt showing the mutation after a certain amount of time. This process is called *monoclonal conversion* and has been observed experimentally for mutations in mitochondrial DNA ^48^ and for the progeny of stem cells ^49^. The latter gives rise to the hypothesis that CRC originates in stem cells, as in these cells the mutations are least susceptible to be washed out of the crypt. The corresponding type of monoclonal conversion starting at the bottom of the crypt is called *bottom-up morphogenesis*.

In contrast, the existence of colonic crypts showing morphologically normal bottom regions paired with APC inactivation at the top and dysplastic epithelium lining the luminal surface ^50^ suggests that mutants at the top of the crypt expand downwards. Additionally, a mathematical model ^51^ predicts that at least the second *APC* hit occurs in the migrating population, as opposed to both hits occurring in stem cells. This type is summarized as *top-down morphogenesis*.

### B | PARAMETER ESTIMATES BASED ON BIOMEDICAL DATA

For simulating the intra-crypt dynamics, we set the parameters of the model according to data published in the literature as well as to educated guesses if no data are available.

#### B.1 | Geometric Initialization of Crypts

As initialization, we set an equidistant grid of 80 × 20 (length × height) nodes and compute the Voronoi tessellation. This leads to a mesh resembling a honeycomb, with symmetrical hexagonal cell shapes. Each cell is then assigned a cell cycle model. As default time step, we set 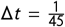 hours.

#### B.2 | TA Cell Cycle and Differentiation Parameters

Estimates of the cell cycle lengths of human cells heavily depend on the type of tissue and the cell type within this tissue. We assume the TA cell cycle to last for approximately 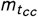 = 24 hours^42^, approximating the duration as follows: The G1 phase lasts about 11 hours, the S phase about 8 hours, the G2 phase about 4 hours, and the M phase about 1 hour. In particular, we assume for the duration of the G1 phase to be normally distributed with 𝒩 (11 hrs, 0.5 hrs).

We assume a general Wnt threshold of *τ*_wnt_ = 0.75 in order to be consistent with existing schematics of the human colonic crypt. Further, according to Equation (5), we assume

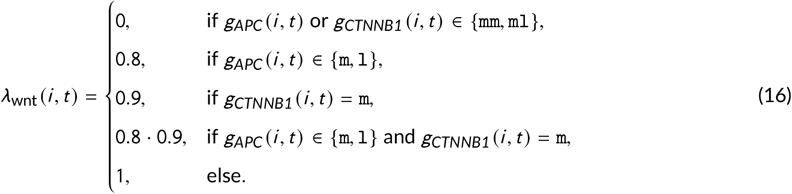

#### B.3 | Mutation Parameters

As depicted earlier, we consider point mutations and LOH events. The derived estimates for point mutation and LOH event rates are in concordance with the ones presented in ^32^.

##### Point Mutations

For modeling the rate *π*_pt_ (gene) of point mutations in a specific gene for each cell, we make the following assumptions:

⊳ In each cell division, we accumulate *n*_pt_ = 10 point mutations.
⊳ The point mutations are uniformly distributed over the base pairs on the entire genome.
⊳ There are *n*_bp,genome_ = 3.2 · 10^9^ base pairs (bp) on the genome.
⊳ Only the point mutations which occur in hotspots of the genes are relevant for the development of a tumor. Hotspots are regions of a gene which give rise to a phenotypical change if mutated. The size of the hotspots *n*_hs_ (gene) is gene dependent and is explained in the following.
⊳ We assume that the alleles are independent of each other, i.e., a mutation in one allele does not influence the mutation probability in the other allele. Thus, the likelihood *π*_pt_ (gene) is twice as large if there is no mutated allele (*n*_mut_ (gene) = 0) compared to the state where one allele is already mutated (*n*_mut_ (gene) = 1).

###### Proposition 2 (Point Mutation Rate)

*Under the assumptions stated above, the rate of a relevant point mutation for a specific gene of a cell is given by*

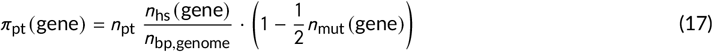

Regarding the hotspots, we assume for *MLH1* and *MSH2* that the whole coding sequence is susceptible to inactivating point mutations, where we use the reference sequence database at NCBI for coding sequence lengths ^52^. For *APC*, we use mutation data from the publicly available DFCI database using the cBioPortal website ^53;54^. We make use of data from about 4000 CRC samples to identify approximately 2400 hotspots. For the present model, we assume for *CTNNB1* that only 5 mutations on 12 base pairs are relevant, according to ^55^. In summary, we obtain the following numbers for *n*_hs_ given in Table 2.

**TABLE 2.** Estimates for the number of hotspots *n*_hs_. The given estimates are used for the computation of the point mutation rates for the individual genes. Those are based on the following data from the literature: *MLH1* and *MSH2*: _^52^_; *CTNNB1*: ^55^; *APC*: ^53;54^.

| Gene | $n_{hs}$ |
| --- | --- |
| *MLH1* | 2,270 |
| *MSH2* | 2,800 |
| *CTNNB1* | 12 |
| *APC* | 2,400 |

##### LOH Events

We assume that all detectable LOH events are large enough to inactivate an affected gene. In other words, we assume that if LOH affects a certain gene, then an exon will be lost and the gene, therefore, is inactivated. As a consequence, the probability of LOH *p*_LOH_ (gene) for a given gene is proportional to its length, denoted by *n*_bp_ (gene).

###### Proposition 3 (LOH Event Rate)

*The rate of a relevant* LOH *event for a specific gene of a cell is given by*

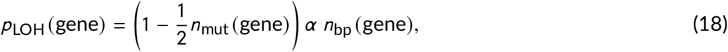

*where α* ∈ ℝ_>0_ *is a parameter to be estimated, independent of the considered gene*.

The available data for *MLH1* suggests that inactivation is twice as likely to occur due to LOH than due to point mutations ^56^. Thus, we assume

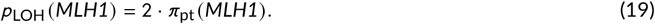

Together with (17) and (18), we get

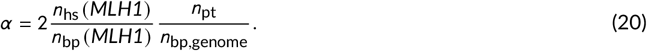

In order to determine *α* and *p*_LOH_, we again use the reference sequence database at NCBI for the length of individual genes ^52^ given in Table 3.

**TABLE 3.** Estimates for the number of base pairs *n*_bp_. The following estimates for *n*_bp_ are necessary for the computation of the LOH rates for the individual genes. They are based on the reference sequence database at NCBI ^52^.

| Gene | $n_{bp}$ |
| --- | --- |
| *MLH1* | 57,500 |
| *MSH2* | 80,000 |
| *CTNNB1* | 41,000 |
| *APC* | 139,000 |

Further, we assume a dependent *CTNNB1* and *MLH1* loss with occurrence rate *r*_effLOH_ = 0.8. In addition, the point mutation rate of MMR-deficient cells is increased by a factor *λ*_mut_ = 100 compared to MMR-proficient cells, since the MMR system, when functioning correctly, only fails to detect 1 out of 100 base pair mismatches (Sec. 7.11 in ^57^).

#### B.4 | Cell Migration Parameters

To the best of our knowledge, no experimental estimates exist for the parameters of the spring force model. We have chosen a parameter combination leading to simulation results which are most consistent with the biological reality. In summary, we set *s* = 1 length unit, *є* = 0.5 length units, *t*_*g*_ = 3 hours, *t* _*a*_ = 0.5 hours, *µ* = 14, *γ* = 0.0675 length units per hour, and *ν* = 1.

#### B.5 | Cell Death Parameters

We assume that all cells with a surface area below *τ*_size_ square length units are not viable. We set the parameter in our simulations

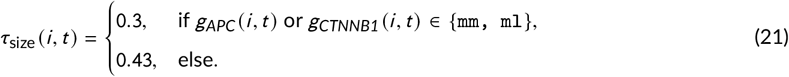

Further, we set the rate of cell death induced by all other homeostatic processes per cell cycle *p*_*cc*_ = 0.0005 and the rate of mutation-induced cell death per cell division *π*_mutdeath_ = 0.0001.

#### B.6 | Stem Cell Parameters

We assume *S* = 6 stem cells per crypt ^6^ with a stem cell cycle duration of ten weeks. The probabilities for stem cell mutations and mutation-induced death are the same as for TA cells. In case of a lethal mutation, an adjacent stem cell divides symmetrically and any of the stem cells populates the crypt. After each stem cell change, the bottom TA cell row is mutated according to the mutations of the newly active cell. The probability of stem cell exchange per stem cell division is set to *p*_change_ = 0.5. Thus, on average, a stem cell will populate the crypt for about 5 months.

### C | PSEUDOCODE

#### Algorithm 1

Pseudocode of the presented computational model.

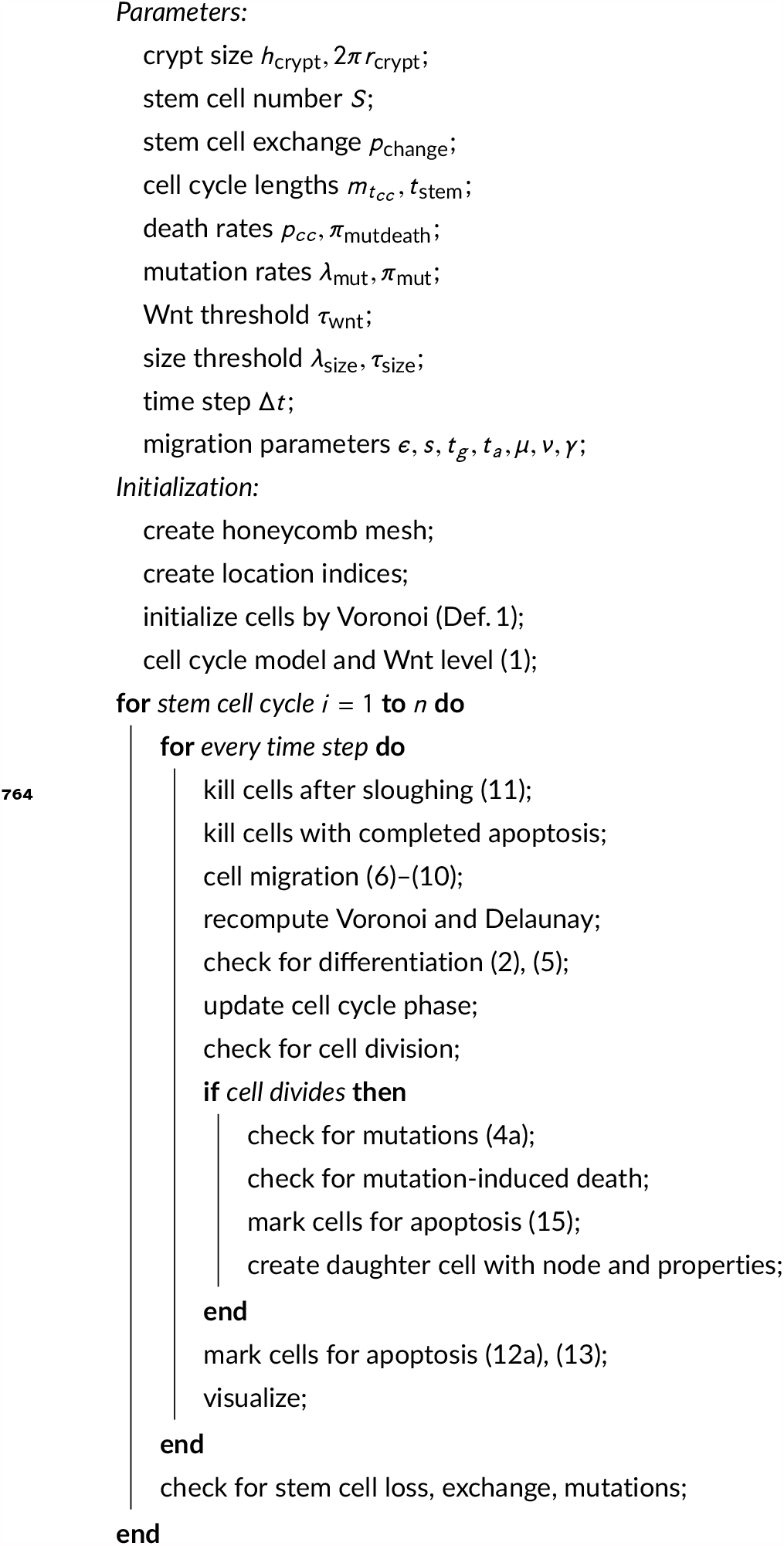

### D | SIMULATION VIDEOS

We provide videos as supporting information with full-time dynamics of the figures in the main text with video frames. The legends of the videos to the corresponding main text figures are given below.

**Video 1 to Figure 9: Loss of MMR deficiency after stem cell exchange**

**Video 2 to Figure 4 (a): Exponential expansion of biallelically *APC*-mutated cells**.

Color legend is as before, biallelically *APC*-mutated cells are shown as dark gray with black nuclei, monoallelically *APC*-mutated cells as light gray. With an MMR-deficient stem cell as initial condition, the second *APC* hit occurred after 20 days in a single-cell monoallelic *APC*-mutated and MMR-deficient clone. Five days after the mutation, a small clone of 32 biallelic *APC*-mutated cells has formed. An additional four days later, the clone has expanded to 400 cells.

**Video 3 to Figure 4 (b): Exponential expansion and monoclonal conversion of biallelically *CTNNB1*-mutated cells**.

**Video 4 to Figure 6 (a): Limited spread of *APC* mutations in an MMR-deficient crypt**.

**Video 5 to Figure 6 (b): Successful spread of an *APC* mutation in an MMR-deficient crypt**.

**Video 6 to Figure 7: Prolonged monoclonal conversion of an MMR stem cell mutation**.

**Video 7 to Figure 8 (a), (b): Top-down morphogenesis of a biallelically *APC*-mutated crypt**.

**Video 8 to Figure 8 (c), (d): Bottom-up morphogenesis of a biallelically *APC*-mutated crypt**.

## Notes

https://github.com/Mathematics-in-Oncology/ComputationalColonicCrypts

### Competing Interest Statement

The authors have declared no competing interest.

